# Tyrosine-based Signals Regulate the assembly of Daple•PARD3 Complex at Cell-cell Junctions during Polarized Planar Cell Migration

**DOI:** 10.1101/717041

**Authors:** Jason Ear, Anokhi Saklecha, Navin Rajapakse, Julie Choi, Majid Ghassemian, Irina Kufareva, Pradipta Ghosh

**Affiliations:** Department of Cellular and Molecular Medicine, University of California San Diego, La Jolla, California 92093; Department of Chemistry and Biochemistry, University of California San Diego, La Jolla, California 92093; Skaggs School of Pharmacy and Pharmaceutical Sciences, University of California San Diego, La Jolla, California 92093; Department of Medicine, University of California San Diego, La Jolla, California 92093; Rebecca and John Moore Comprehensive Cancer Center, University of California San Diego, La Jolla, California 92093; Veterans Affairs Medical Center, La Jolla, CA

## Abstract

Polarized distribution of organelles and molecules inside a cell is vital for a range of cellular processes and its loss is frequently encountered in disease. Polarization during planar cell migration is a special condition in which cellular orientation is triggered by cell-cell contact. Here, we demonstrate that the multi-modular signaling scaffold Daple (CCDC88C) is a component of cell junctions in epithelial cells which serves like a cellular ‘compass’ for establishing and maintaining contact-triggered planar polarity *via* its interaction with the polarity regulator PARD3, which has been implicated in both apical-basal and planar polarity. This interaction, mediated by Daple’s PDZ-binding motif (PBM) and the third PDZ domain of PARD3, is fine-tuned by two tyrosine phosphoevents on Daple’s PBM that are known to be triggered by a multitude of receptor and non-receptor tyrosine kinases, such as Src. Hypophosphorylation strengthens the interaction, whereas hyperphosphorylation disrupts it. These findings reveal an unexpected role of Daple within the planar cell polarity pathway as a platform for signal integration and gradient sensing for tyrosine-based signals.

## Introduction

Epithelial cell-cell adhesion and cell polarity are necessary for proper cell function and tissue organization [1]. Cells along an epithelium are connected together through various junctional complexes (tight junction, adherens junctions, desmosomes, and gap junctions) which are essential in establishing the apical and basolateral pole of a cell (referred to as the apical-basal axis) [2]. In addition to an apical-basal polarity, cells are also polarized along the plane of the epithelium, referred to as planar cell polarity (PCP) [3]. Dysregulation of these cell adhesion complexes and cell polarity can lead to maladies such as inflammatory bowel diseases, hydrocephalus, abnormal skin barrier function and tumor initiation and progression [4]. In fact, disruption of cellular junctions is one of the first events during epithelial-to-mesenchymal transition (EMT), a phenomenon that is encountered during cancer initiation and progression [4, 5]. Thus, cell-cell junctions are widely believed to serve as tumor suppressors [6, 7].

A typical cell junction is comprised of a layer of transmembrane molecules which functions for hetero- and homotypic binding between cells, and a layer of molecular scaffolds peripheral to the membrane. These scaffolds tether transmembrane molecules to key intracellular molecules such as the cytoskeleton or signaling molecules. Many of these scaffolds contain one or more PDZ domains [<u>P</u>ost synaptic density protein (PSD95), <u>D</u>rosophila disc large tumor suppressor (Dlg1), and Zonula occludens-1 protein (zo-1)], which mediate interactions with other proteins that contain a PDZ-binding motifs (PBM) [8]. Mutations affecting PDZ•PBM interactions have exposed the importance of these junction-localized interactions in the regulation of key tumor cell phenotypes [8].

Daple (CCDC88C) is a large, multi-modular PBM-containing scaffold protein [9, 10]. First discovered in a yeast-two hybrid screen through its ability to bind to the PDZ domain on Dishevelled (Dvl), Daple has emerged as a key modulator of Wnt signaling [9, 11]. The extreme C-terminus on Daple contains an ‘atypical’ PBM which we, and others, demonstrated to be necessary for the Daple•Dvl interaction [10–12]. We subsequently showed that Daple binds Gαi proteins and triggers GTPase signaling downstream of the Wnt receptor Frizzled-7 upon activation by Wnt5a; such signaling is necessary for the activation of the β-catenin-independent Wnt signaling pathway (a.k.a. non-canonical Wnt signaling). More recently, we showed Daple-dependent Wnt signaling is also shaped by Akt and multiple tyrosine kinases (TKs); in each case, phosphorylation of Daple by these kinases impacted its localization and signaling [9, 10, 13]. We also showed that Daple has two opposing roles during colorectal cancer (CRC) progression; much like the non-canonical Wnt pathway, Daple acts as a tumor suppressor in the normal epithelium and early during cancer initiation, but it serves as a potent trigger for EMT and tumor cell invasion later during cancer dissemination. How does one protein serve two seemingly opposing roles, and how its tumor suppressive and pro-metastatic functions are segregated remains poorly understood.

Here we show that Daple’s functions are tightly regulated by spatial segregation, which is dictated by its interactome and the context, i.e., cell type and key phosphomodifications that are triggered by growth factors. Such fine-tuned regulation of Daple’s subcellular localization and its interactome offers insight into Daple’s dual role during cancer progression and other cell polarity associated diseases.

## Results and Discussion

### Daple is localized on cell-cell junctions in epithelial cells and tissues

To investigate the role of Daple in the normal colonic epithelium, we first studied the expression and subcellular localization of Daple using a previously validated antibody against its C-terminus in a variety of immunocytochemical approaches. By immunohistochemistry (IHC) on formalin-fixed paraffin-embedded (FFPE) colon biopsies, we confirmed that Daple is indeed expressed in the epithelial lining of the human colon (**Figure 1A**). By immunofluorescence staining and confocal imaging of human colon-derived organoids cultured in matrigel, we determined the localization of Daple at a higher resolution; it localizes at the apical side of the epithelial cells facing the lumen, marked by the tight junction protein, occludin. Findings suggest that Daple is on the apical pole of the epithelial cells (**Figure 1B**). By immunofluorescence on monolayers of cultured cancer cell lines, we observed Daple at the sites of cell-cell contact (**Figure 1C**). In the highly polarized MDCK cells, we observe colocalization with the tight junction marker occludin as well as with the adherens junction marker, β-catenin (**Figure 1D**). Taken together, these findings demonstrate that a significant pool of the cytosolic protein Daple localizes to cell-cell junctions. This is in keeping with prior observations made by others using ependymal, inner ear and MDCK cells [13–17]. These findings suggest that Daple may physically associate with various junctional complexes to maintain such localization.

**Figure 1.**
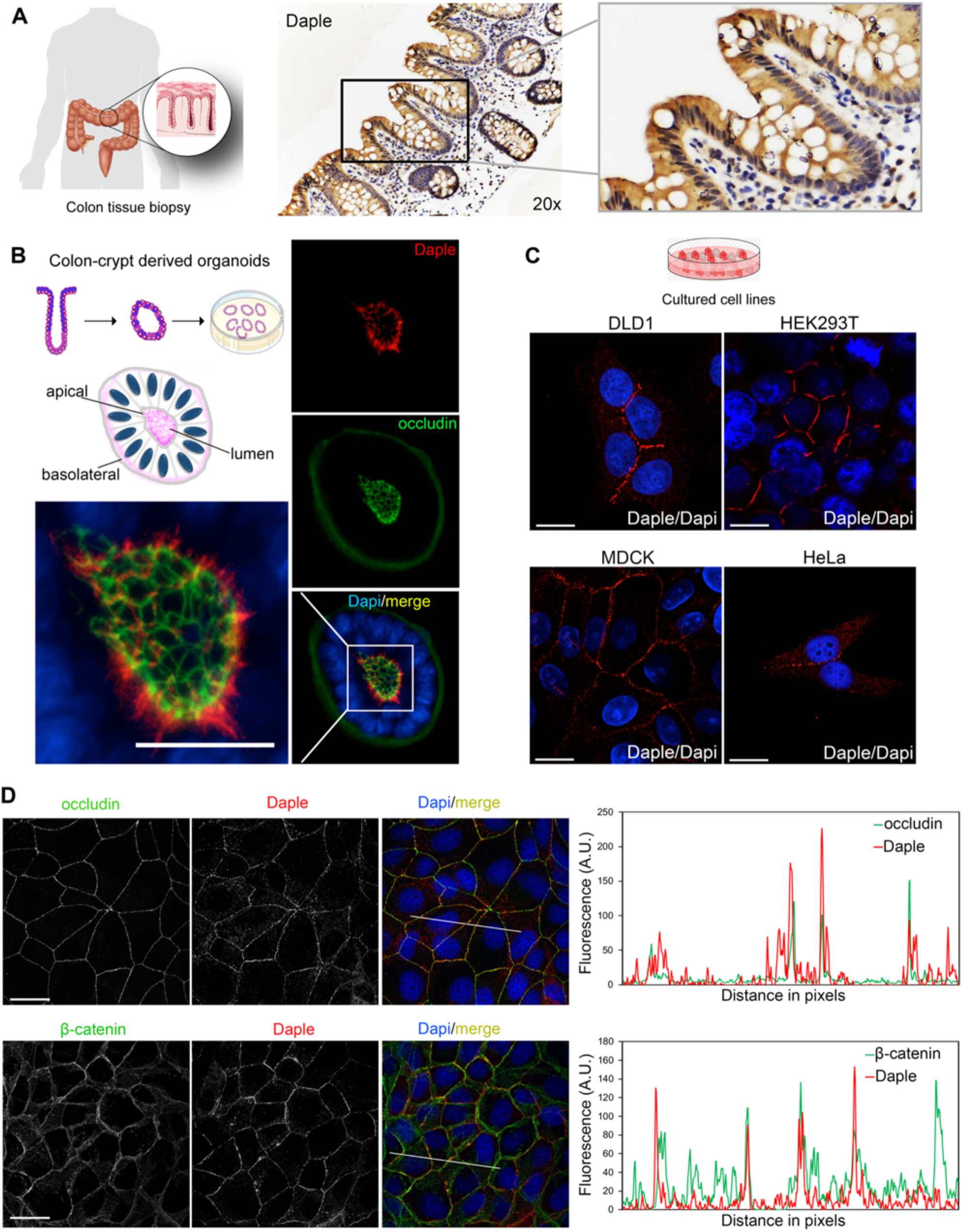
Daple localizes to the cell-cell junctions in polarized epithelial cells. **A)** Immunohistochemistry staining on human colon biopsy sections showed that Daple is highly expressed in the colon epithelium. **B)** Human colon-derived organoids were fixed and stained for Daple (*red*) and occludin (*green*) and analyzed by confocal microscopy. A representative image is shown; scale bar, 25 μm. **C)** Various epithelial cell lines (DLD1, HEK293T, MDCK, and HeLa) were fixed, stained for Daple (red) and nucleus (Dapi) and analyzed by confocal microscopy. Scale bar, 10μm. **D)** MDCK cells were fixed, stained for Daple (*red*) and either the tight junction marker, occludin (*green, top*) or the adherens junction maker, β-catenin (*green, bottom*) and analyzed by confocal imaging. Scale bar, 10μm. *Left*: Representative images are shown. *Right*: RGB profile plot of indicated region is shown.

### Daple localizes to cell junctions in well-differentiated CRC cells, requires junctional complexes

Next, we examined the subcellular localization of Daple in several CRCs cell lines. Daple was found at cell-cell junctions in the poorly metastatic CaCo-2 cells but was predominantly cytosolic in the highly metastatic Sw480 cells (**Figure S1A**). The total levels of expression of Daple in both cell lines are comparable (**Figure 2A**), leading us to conclude that the observed differences in localization are likely due to the absence of cell junctions in Sw480 cells.

**Figure 2.**
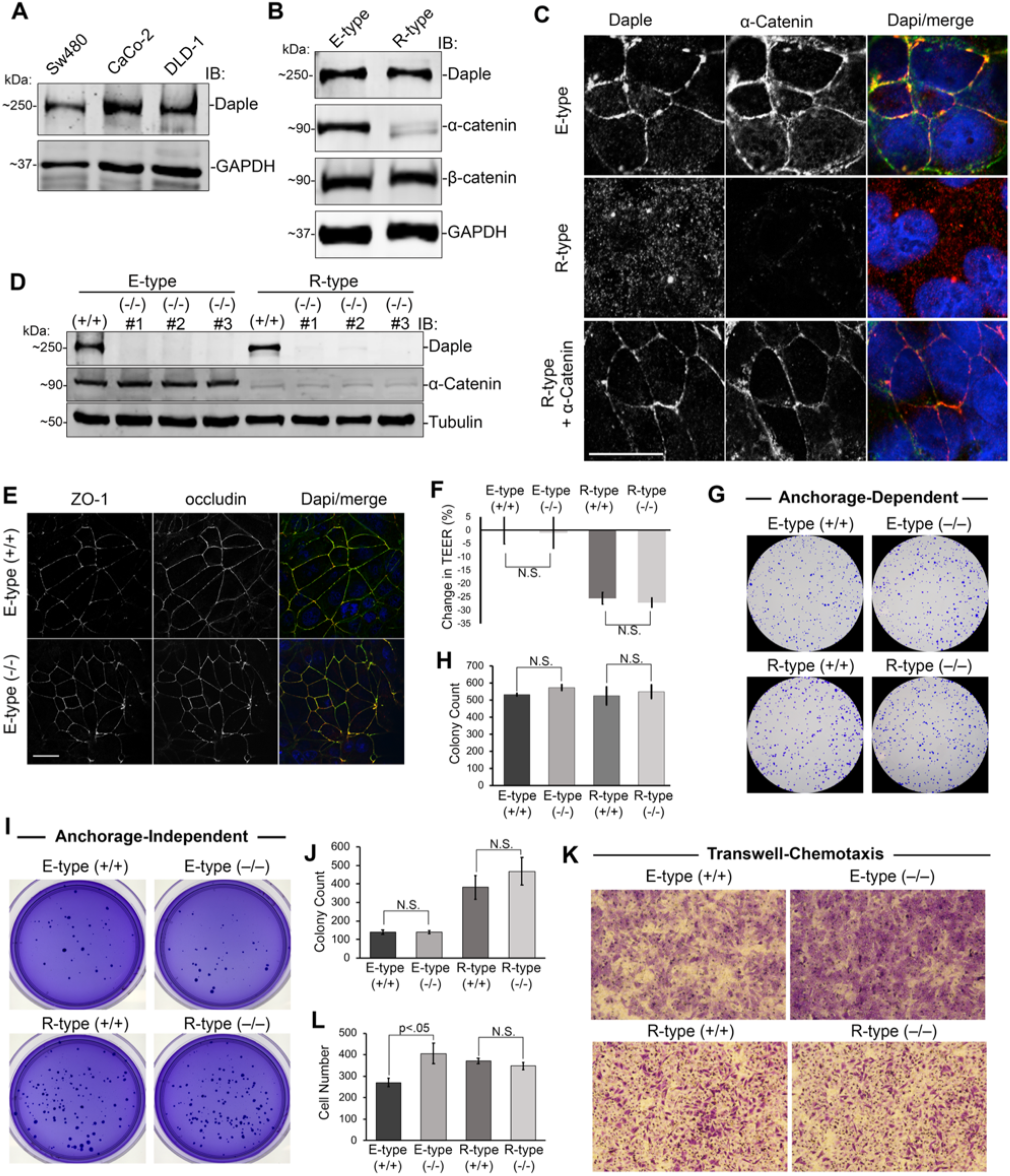
Depletion of Daple in the poorly migrating DLD1 E-type cells vs. the highly invasive R-type cells have differential effect on cell migration. **A)** Immunoblot of whole cell lysates of Sw480, CaCo-2, and DLD1 colorectal cancer cells for Daple expression. **B)** Immunoblot of whole cell lysates of DLD1 E-type and R-type cells for Daple and α-catenin expression. **C)** DLD1 E-type, R-type, and R-type cells transiently expressing α-catenin were fixed and immunostained for Daple (*red*) and α-catenin (*green*). Scale bar, 5μm. **D)** Immunoblot of DLD1 E-type and R-type CRISPR/Cas9 clones targeted for Daple confirming depletion. **E)** DLD1 E-type cells depleted of Daple (−/−) were stained for cell junction markers ZO-1 (red) and occludin (*green*); representative images are shown. Scale bar, 25μm. **F)** Transepithelial electrical resistance (TEER) of DLD1 E-type and R-type cells depleted (−/−) or not (+/+) of Daple. Measurements are represented as percent change relative to E-type (+/+) cells. **G-H)** Anchorage-dependent colony growth assay on DLD1 E-type and R-type depleted (−/−) or not (+/+) of Daple. Bar graphs (H) show quantification of panels in G. **I-J)** Anchorage-independent growth assay on DLD1 E-type and R-type cells depleted (−/−) or not (+/+) of Daple. Bar graphs (J) show quantification of panels in I. **K-L)** Transwell migration assay on DLD1 E-type and R-type cells towards 2% serum. Bar graphs (L) show quantification of panels in K.

To further investigate this differential localization, we took advantage of the fact that most CRC cell lines have a mix of well-differentiated non-invasive epithelioid (henceforth, E-type) and poorly-differentiated highly invasive rounded (henceforth, R-type) cells representing transition states of CRC progression [18]. We studied two well-characterized DLD1 E-type and R-type lines [19]; the latter lacks the adherens junction protein, α-catenin. We chose to study this pair because loss of α-catenin has previously been shown to be associated with a poorly differentiated status of CRCs and a worse prognosis in patients (**Figure S1B**). Several clones of E-type and R-type cells (see **Figure S1C** for distinct morphology) were isolated and the loss of α-catenin in R-type cells was confirmed by immunoblotting (**Figure S1D**). Although both clones appear to express similar levels of Daple protein, as determined by immunoblotting (**Figure 2B**), localization of Daple in these cells looked as different as that previously observed between CaCo-2 and Sw480; Daple was at cell-cell junctions in the E-type cells, but in the cytoplasm in R-type cells (**Figure 2C**). It is noteworthy that many components of cell junctions are insoluble in detergents such as triton x-100 (Tx100) due to their association with the cortical actin at the cell periphery. Daple was detected in Tx100-insoluble fractions in all cell types tested (**Figure S2A-C**), suggesting that regardless of its junctional localization, Daple remains associated with the actin cytoskeleton.

Next, we carried out a series of studies to dissect whether Daple’s localization to cell-cell junctions is an active process of recruitment to that site or a passive consequence due to the formation of these junctions. Transient expression of α-catenin into R-type cells restored the localization of Daple at cell junctions, mimicking E-type cells (**Figure 2C**). Conversely, disruption of cell junctions in E-type cells by calcium depletion (using the calcium-chelator, EGTA) triggered redistribution of Daple from cell junctions to the cytoplasm (**Figure S1E**). Prior work showed that activation of PKC in R-type cells (using TPA, tetradecanoyl phorbol acetate) transiently restores junctions [20]; although we too observed an increase in the localization of occludin at the junctions upon TPA treatment, Daple did not localize there (**Figure S1F**). These findings demonstrate that the assembly and disassembly of adherens junctions is a key determinant of Daple’s localization at cell junctions. The observation that exogenous expression of α-catenin, but not TPA-treatment, was sufficient to restore Daple’s localization at cell junctions suggests that Daple’s ability to localize to this subcellular site is an active process of recruitment which may require interactions with stable junctional structures.

### Junction-localized Daple enables contact-triggered orientation and polarized planar migration

To investigate if junction-localized Daple may regulate key cellular phenotypes, we depleted Daple in the E-type and R-type cells by CRISPR/Cas9, and confirmed by immunoblotting (**Figure 2D**, **S3A-C**). Depletion of Daple in E-type did not alter the levels of α-Catenin (**Figure 2D**) and did not have a discernible impact on the morphology of cell junctions, as evident by localization of the integral membrane protein occludin and the peripheral tight junction protein ZO-1 (**Figure 2E**). There was also no discernible impact on the functional integrity of tight junctions, i.e., regardless of Daple depletion, the paracellular permeability remained unchanged, as determined by measurements of transepithelial electrical resistance (TEER) on a confluent monolayer of cells (**Figure 2F**). These findings indicate that loss of Daple may be dispensable for the morphological or functional integrity of tight junctions in epithelial cells grown in confluent monolayers. Findings are also consistent with what has been reported by others for tight junctions in the blood brain barrier or Daple −/− mice [15].

Because junctions are also known to serve as signaling microdomains that dictate tumor-sphere growth and cell migration [4–7, 21], next we asked if these growth and motility phenotypes are impacted in Daple-depleted cells. Under the conditions tested, we observed no significant changes in growth either in anchorage-dependent or in anchorage-independent colony formation assays (**Figure 2G-H, I-J**). When we measured the ability of these cells to migrate across a semipermeable membrane across a 0-to-2% serum gradient, loss of Daple led to an increase in chemotaxis in the E-type but not in the R-type cells (**Figure 2K-L**), indicating that loss of junctional Daple induces gradient-responsive migration in 3-dimension (3D).

Because Daple is an enhancer of non-canonical Wnt signaling [9, 11, 12], signals that mediate the phenomenon of orderly polarized migration over 2D planes, a.k.a planar cell polarity (PCP) [3, 15], we asked if junction-localized Daple may impact this phenomenon. PCP is a critical tumor suppressive phenomenon that maintains homeostasis in the colon crypt by ensuring that the cells newly generated from the crypt base-localized stem cells migrate along the basement membrane (2D migration on a plane) to constantly replenish the crypt top-localized terminally differentiated surface epithelial layer which displays a high turnover rate [22, 23]. When studying PCP in cell culture, migrating cells are typically monitored on a planar surface [24, 25]. While this mode of migration is more accurately accessing front-rear polarity, many components in front-rear polarity is conserved in PCP [24]. We utilized one such previously described planar cell migration model [26] in which cells are first grown in suspension to form a compact spheroid, then transferred onto an adherent substrate and monitored by light and confocal microscopy for polarized radial migration (**Figure 3A**). We chose fibronectin as the substrate because it is well-accepted as the matrix of choice for PCP studies [27, 28]. While no differences were observed between spheroids grown using wild-type and Daple knockout E-type cells (**Figure S3D and E**), Daple depleted cells migrated out significantly more than wild-type cells, losing cell-cell contacts (higher birefringence in light microscopy, **Figure 3B**) covering a greater surface area (**Figure 3B and C**). These migrating cells also displayed a loss of cell-cell contact (marked by the adherens junction marker, E cadherin) and loss of contact-triggered orientation towards the spheroid, and instead polarized towards the free-edge, as determined by the relative positions of the nucleus and the Golgi body (marked by the integral membrane protein, GM130) [29, 30]. Compared to wild-type cells, a greater number of Daple-depleted cells positioned their Golgi away from the spheroid, indicating that cells were polarized towards the free-edge, which is consistent with their observed increased motility in both 2D and 3D assays (**Figure 3D and E**). Overall, these finding implicate Daple in front-rear polarity (and presumably PCP); its loss disrupted contact-triggered orientation and migration, and instead promoted contact-free scattering. Because others have implicated Daple in regulating apical-basal polarity and apical constriction in epithelial cells [17], taken together, these findings indicate that junction-localized Daple may serve as a nexus between two types of polarity—apical-basal’ and ‘front-rear’. Because Daple serves this role in the context of sheet-like migration of cells on a planar surface, findings suggest it regulates PCP.

**Figure 3.**
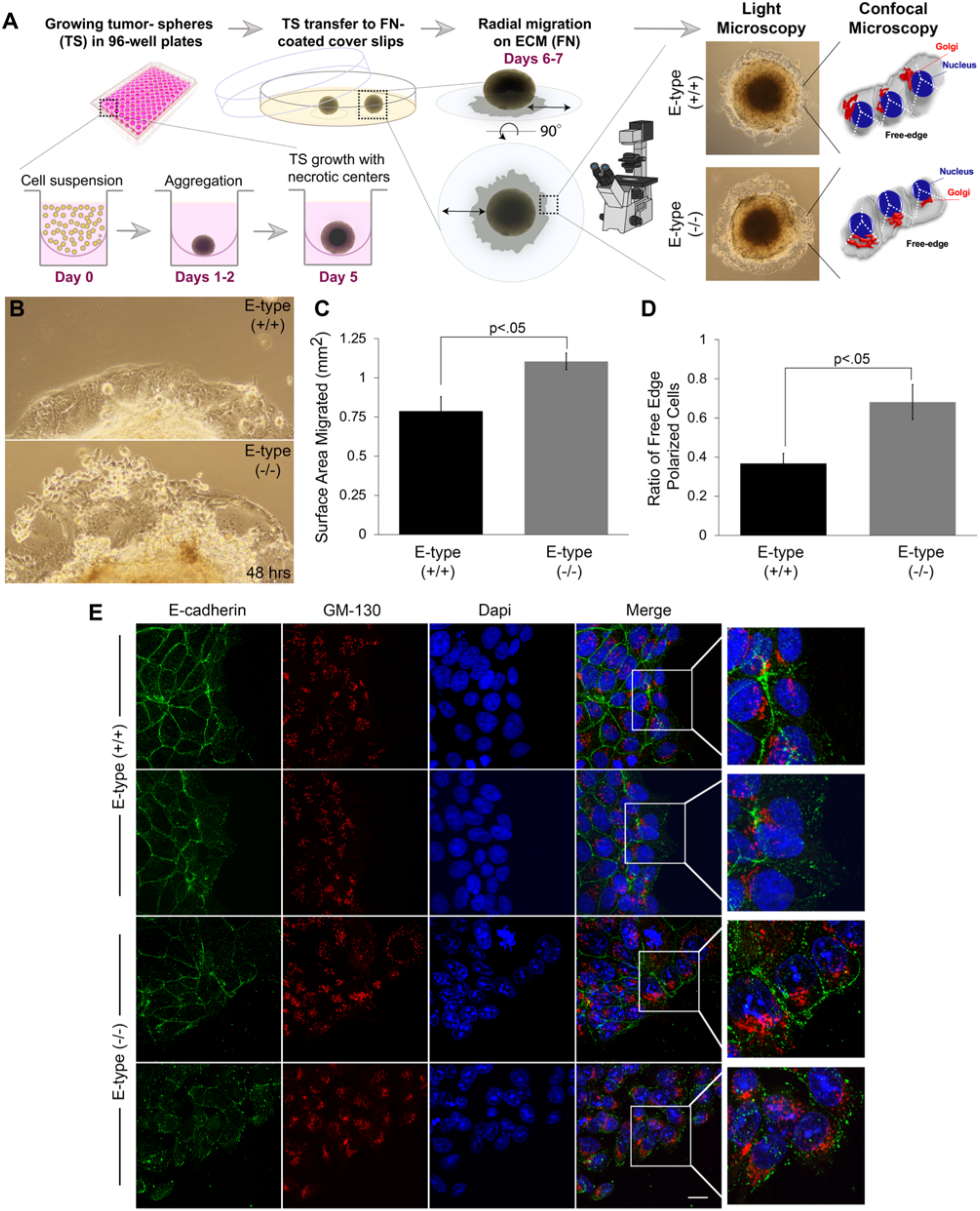
Loss of Daple affects planar migration of DLD1 E-type cells. **A)** Schematic of workflow for tumor spheroid growth and assessment of radial migration by light and confocal microscopy. Polarization of migrating cells was monitored by assessing the position of the Golgi (using GM-130) relative to the advancing edge and the nucleus (using DAPI). **B)** Magnified view of the edge of the tumor sphere showing radially migrating sheets of E-type (+/+) and (−/−) cells after 48 hours, as visualized by light microscopy. **C)** Bar graphs show the extent of surface area of sheet-like migration in B. **D-E)** Radially migrated cells, as in C, were fixed and stained for the *bona-fide* Golgi marker GM130 (*red*) and E-cadherin (*green*). The ratio of polarized cells to total cells was quantified (E).

### Biotin proximity labeling identifies various PDZ-proteins within Daple’s interactome

We hypothesized that junction-localized Daple may impact PCP via the coordinated regulation of signaling co-complexes that sense and respond to cell-cell contact via adherens junctions (but may not regulate tight junction integrity). To gain insights into how/why Daple localizes to the cell junctions and impacts PCP, we carried out BioID proximity labeling coupled with mass spectrometry (MS) to identify interacting proteins. We carried out these studies in both in E- and R-type cells and HEK293T cells in order to specifically understand which interactions are specific for junction-localized Daple. To this end, Daple was N-terminally tagged with BirA biotin ligase (**Figure S4A**), and the construct was rigorously validated using several approaches. Immunofluorescence studies confirmed that the exogenously expressed tagged construct localized similarly to the endogenous protein—it is found on cell junctions and the perinuclear recycling compartment (**Figure S4B-C**) as described previously [13]. Biotinylation *in situ* was confirmed by incubating cell lysates with streptavidin beads and blotting using fluorescent conjugated streptavidin (**Figure 4A and B**). Immunoblotting and biochemical interaction assays confirmed that the construct was expressed as full-length protein and retained binding to Dvl (**Figure S4D and E**). Staining for biotinylated protein in HEK293T cells revealed that the construct can indeed label proteins at cell junctions (**Figure 4B**).

**Figure 4.**
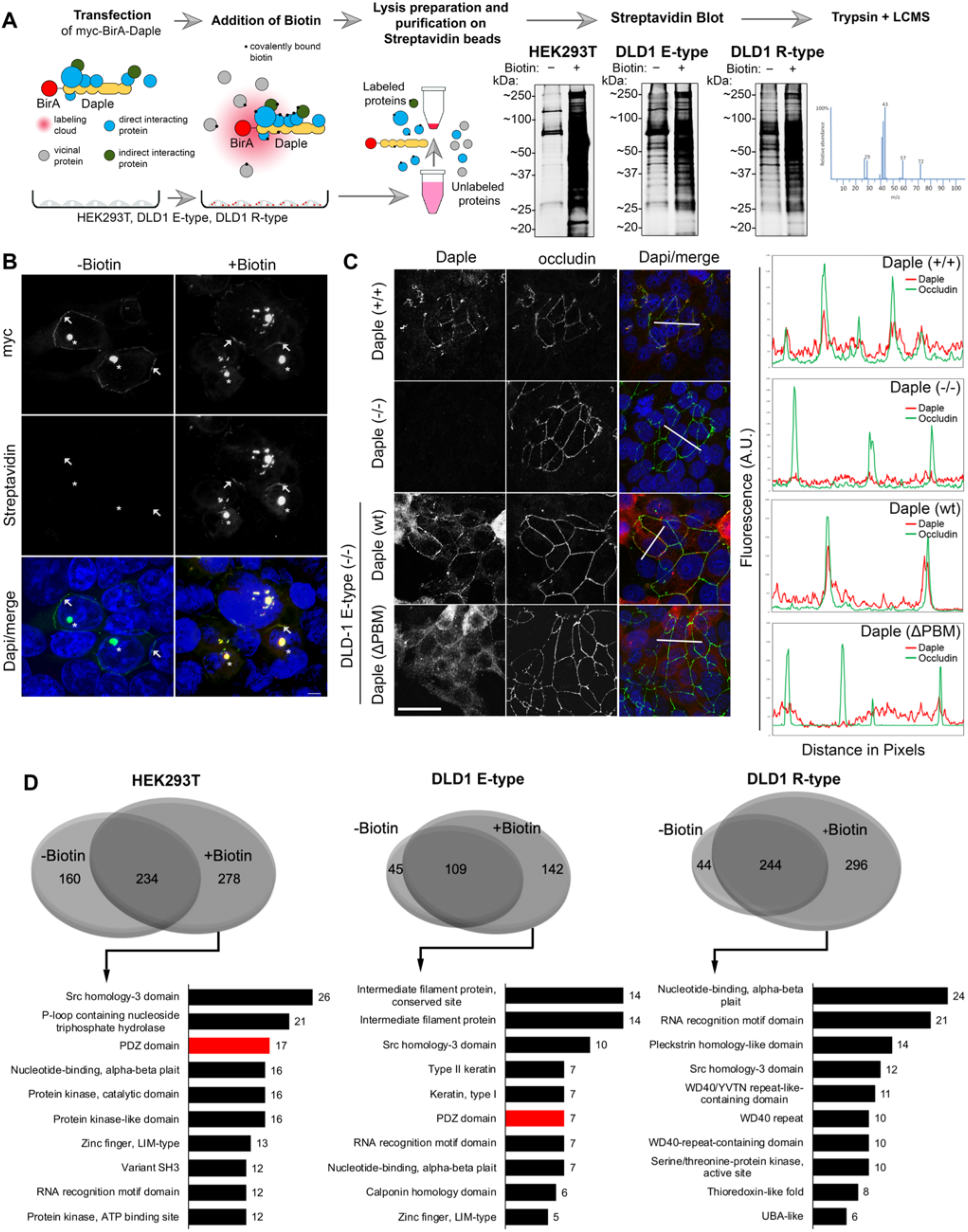
Biotin proximity labeling identifies an enrichment of PDZ proteins within Daple’s interactome in E-type, but not in R-type cells. **A)** Schematic summarizing the workflow of biotin proximity labeling study carried out using exogenously expressed myc-BirA-Daple in various cell lines. Immunoblot with AlexaFluor-680 conjugated streptavidin confirmed biotinylation of affinity purified proteins. **B)** HEK293T cells transfected with myc-BirA-Daple and treated with biotin were stained with AlexaFluor-594 conjugated streptavidin and antibodies against the myc-tag (*green*). Asterisk = pericentriolar localization; arrow = PM localization. Scale bar, 5 μm **C)** DLD1 E-type (+/+), E-type (−/−), or Daple (−/−) cells ectopically expressing Daple-WT or Daple-PBM deficient mutants (ΔPBM) were stained for Daple (*red*) and occludin (*green*). <u>*Right*</u>: RGB plot of region indicated in merge panels. **D)** <u>*Top*</u>: Venn Diagram showing overlap of affinity-captured proteins identified by mass spectrometry between biotin treated or no biotin treated control samples. <u>*Bottom*</u>: Bar graph summarizing identified proteins (exclusive to plus biotin conditions) grouped by protein domain using DAVID GO analysis. Top domains categories are shown.

MS identification of biotinylated proteins in HEK293T, E-type, and R-type cells revealed several novel binding partners of Daple (**Figure 4D**). Because Daple’s C-terminal PDZ-binding motif (PBM) was deemed essential for its localization to cell junction (**Figure 4C**), we carried out gene-ontology (GO) analyses on the MS hits by protein domains (using DAVID GO and INTERPRO domains, with p value <.05 set as significant) expecting to find PDZ-containing proteins that are enriched or eliminated depending on Daple’s localization (**Figure 4D**). Indeed, we found that several PDZ proteins that are associated with cell junctions were enriched in the BioIDs from HEK293T and E-type cells, but not in R-type cells (**Figure 4D and 5A**). It is noteworthy to point out that although R-type cells did not present a significant enrichment for PDZ proteins, two proteins containing PDZ domains were identified (**Figure 5A**). These findings led us to hypothesize that Daple may be recruited onto cell junctions by its PBM module via its interaction with PDZ proteins. Henceforth, we directed our efforts at validating *in vitro* some of the putative PDZ-PBM interactions and understanding how those interactions may be reversibly regulated to allow context dependent localization of Daple at cell junctions.

**Figure 5.**
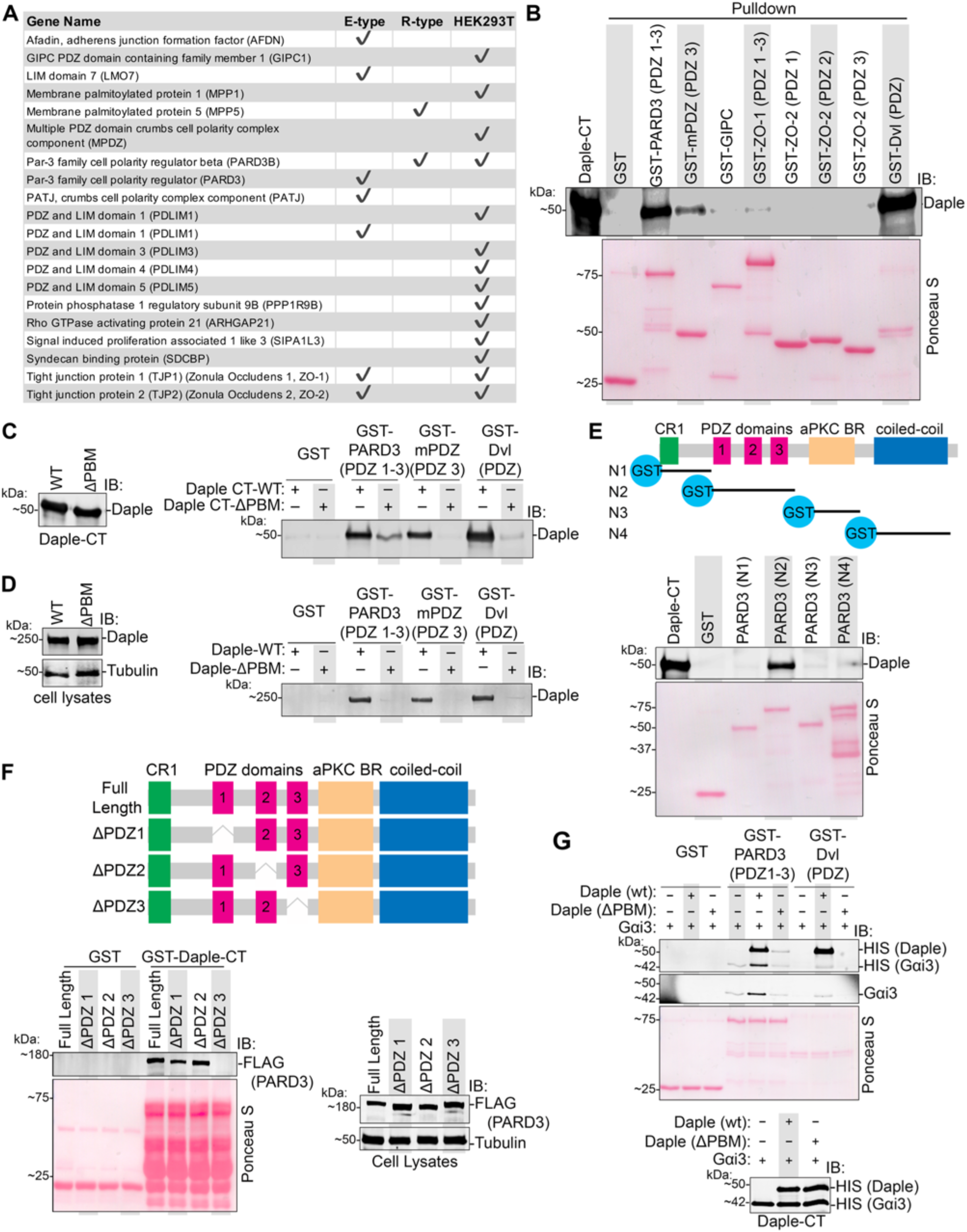
Daple directly and selectively binds to the third PDZ module of PARD3 via its C-terminal PBM and can form ternary co-complex with Gαi3. **A)** Table of PDZ proteins identified by mass spectrometry in BioID studies in Fig 4. **B)** Pulldown assays using purified GST-tagged PDZ domains PARD3, mPDZ, ZO-1, ZO-2, and Dvl immobilized on glutathione beads and soluble recombinant His-Daple-CT. Bound Daple-CT was determined by immunoblotting. **C)** Pulldown assays using GST-tagged PARD3, mPDZ, or Dvl PDZ domains used in binding assays with purified His-Daple-CT-WT or His-Daple-CT-ΔPBM. **D)** Pulldown assays using GST-tagged proteins, as above, where lysates of transiently transfected HEK293T cells was used as source of full length Daple-WT or Daple-ΔPBM. **E)** <u>*Top*</u>: Schematic shows the modularity of various GST-PARD3 constructs used (N1-N4). <u>*Bottom*</u>: Pulldown assays using various GST-PARD3 regions with Daple-CT. GST-PARD3 bound Daple-CT was analyzed by immunoblotting. **F)** Schematic shows the modularity of various FLAG-PARD3 constructs exogenously expressed in cells and used as source of PARD3 for pulldown assays. <u>*Bottom*</u>: Pulldown assays using various FLAG-tagged PARD3 constructs with recombinant GST-Daple-CT. Daple-CT-bound PARD3 proteins (left) and input lysates for FLAG-PARD3 (right) were analyzed by immunoblotting (Left and right, respectively). **G)** Pulldown assays with GST-tagged PARD3 or Dvl PDZ proteins immobilized on glutathione beads and various His-tagged proteins, either alone or in combination, as indicated. Bound complexes and inputs were analyzed by immunoblotting.

### Daple directly and specifically binds to PDZ proteins, PARD3 and mPDZ

BioID relies on transient transfection; hence, it is prone to artifacts due to either compromised cell health or higher than physiologic levels of proteins with resultant altered stoichiometry, and only indicates proximity, not direct interaction. We sought to validate our BioID ‘hits’ by asking which PDZ proteins identified by BioID (**Figure 5A**) directly interact with Daple using recombinant GST-tagged PDZ proteins in *in vitro* interaction assays with purified Daple-CT (aa. 1650 to 2028). Five proteins were prioritized based on the criteria that they are all *bona-fide* PDZ family of proteins that localize to cell junctions: (i) the Par-3 Family Cell Polarity Regulator PARD3; (ii) the Multiple PDZ Domain Crumbs Cell Polarity Complex Component mPDZ; (iii) the tight junction protein 1, TJP1, a.k.a, zonula occludens (ZO)-1, (iv) the tight junction protein 2, TJP2, a.k.a, zonula occludens (ZO)-2, and (v) GAIP-interacting protein, C terminus (GIPC) [31–34]. Interaction was detected with PARD3 and mPDZ, besides the previously known interacting partner, Dvl. A much weaker interaction was detected between ZO-1 and Daple (**Figure 5B**). As suspected, all these interactions were virtually lost when we used purified Daple-CT lacking the described C-terminal PBM (ΔPBM) (**Figure 5C**) or when cell lysates of exogenously expressed full-length Daple (WT or ΔPBM) was used in the interaction assays (**Figure 5D)**. These findings confirm that Daple’s interactions with diverse PDZ proteins are likely to be mediated via a PDZ•PBM interaction.

Because both PARD3 and mPDZ are molecular scaffolds that have more than one PDZ domain, we asked if the binding of Daple to these proteins is mediated specifically via one or more of these domains. When we purified each of the 13 PDZ domains of mPDZ from bacteria as GST-tagged proteins and used them in pulldown assays, we found that Daple preferentially bound the third PDZ domain on mPDZ (**Figure S5A and B**). We took a slightly different approach in the case of PARD3, which contains three PDZ domains (**Figure 5E**). First, we confirmed that binding between Daple and PARD3 is specific to PARD3’s PDZ domains [31], via in vitro protein interaction assays with various GST-tagged PARD3 truncation constructs and recombinant His-Daple-CT. Daple specifically binds to the PARD3 construct which contains the PDZ domains (**Figure 5E**). Next we determined which of the three PDZ domains Daple binds to, by investigating GST-tagged Daple-CT interaction with PARD3 constructs from which individual PDZ domains were deleted [35]. Only deletion of the third PDZ (PDZ3) on PARD3 led to loss of binding to Daple (**Figure 5F**). Taken together with our prior in vitro binding studies, we conclude that Daple directly binds to both PARD3 and mPDZ. In each case, the PBM on Daple is required, and the interaction occurs via the ability of the PBM to bind the third PDZ domains of PARD3 or mPDZ. In doing so, Daple’s appears to display a high degree of specificity because it binds to only one of 13 PDZ modules on mPDZ and one of three modules in PARD3 (**Figure 5F and S5**). Thus, the Daple•PBM recognition is specific to a limited set of PDZ proteins and domains.

### Daple, PARD3 and G protein can form ternary complexes

Prior work in Drosophila has demonstrated the importance of PARD3 and Gαi in regulating cell polarity and centrosome positioning [36, 37]. Because Daple binds and modulates Gαi activity [9], and we now find that it binds PARD3 and regulates PCP, we hypothesized that the Daple•PARD3 interaction may allow scaffolding of G proteins to PARD3, and hence coordinate the establishment of PCP. We began by asking if the newly discovered Daple•PARD3-PDZ interaction affects the Daple•Gαi interaction. We had previously described an unexpected allosteric phenomenon in which the Daple•Dvl-PDZ and the Daple•Gαi interactions are mutually exclusive, i.e., ternary complexes between all 3 proteins, Gαi•Daple•Dvl is not feasible [10]. To our surprise, such allostery was not observed in the case of PARD3; instead, we found that Gαi3 bound PARD3 exclusively in the presence of Daple, and not directly, indicative of the formation of tertiary Gαi•Daple•PARD3 complexes (**Figure 5G**). This suggest that despite both interactions (Daple•Dvl and Daple•PARD3) occurring *via* the same Daple’s PBM, the exact mechanism of binding may differ between various PDZ domains. Moreover, context-dependent interaction of Daple’s PBM with one (Dvl) or the other (PARD3) may impede or enable, respectively, Daple’s ability to bind and modulate G protein signaling.

### Tyrosine phosphorylation of Daple’s PBM shapes its PDZ-interactome, localization to cell junctions

The two tyrosines immediately adjacent to Daple’s PBM have previously been shown to serve as a platform for convergence of signaling downstream of multiple receptor and non-receptor tyrosine kinases (TKs) like Src [10] (**Figure 6A**). We asked if these phosphoevents can dictate which PDZ proteins bind Daple. As shown previously, tyrosine phosphorylation of Daple’s PBM by Src reduced binding of Daple to Dvl [10]; by contrast, phosphorylation enhanced binding to PARD3 under the same conditions (**Figure 6B and C**). We dissected the relative contributions of single phosphorylations at either Y2023 or Y2025 using GST-tagged Daple-PBM (aa 2008-2028) and a previously published strategy of using phosphomimicking mutants [Tyr(Y) to Glu(E)] [10]. Mutating either tyrosine residue to glutamate alone did not decrease binding to PARD3, instead, Y2023E enhanced binding to PARD3 (**Figure 6D-E**). By contrast, mutating either Y to E was sufficient to disrupt binding to Dvl (**Figure 6D-E**). The dual phosphomimicking Daple construct bound neither PARD3, nor Dvl. In converse experiments, where GST-PARD3 or GST-Dvl is used to pulldown His-Daple-CT, we see similar findings, in that Daple Y2023E alone was sufficient to disrupt the Daple•Dvl interaction (consistent with prior report [10]), but not the Daple•PARD3 interaction (**Figure S6A-C**). Findings from GST-pulldown assays were recapitulated in coimmunoprecipitation studies in cells; immune complexes showed that PARD3 interacts with ectopically expressed Daple-WT or Daple-FA (F1675A, a previously described G-protein binding deficient mutant) but not with Daple-ΔPBM or the dual phosphomimick Daple-Y2023/2025E mutant (**Figure 6F**). This dual phosphomimic mutant failed to localize to cell junctions (**Figure 6G**), which is consistent with the defect we observed earlier in the case of the Daple-ΔPBM mutant (**Figure 4C**). Finally, cells retain the ability to localize PARD3 onto junctions in the absence of Daple (**Figure S6D**), suggesting that Daple is recruited onto junctions by PDZ domain proteins.

**Figure 6.**
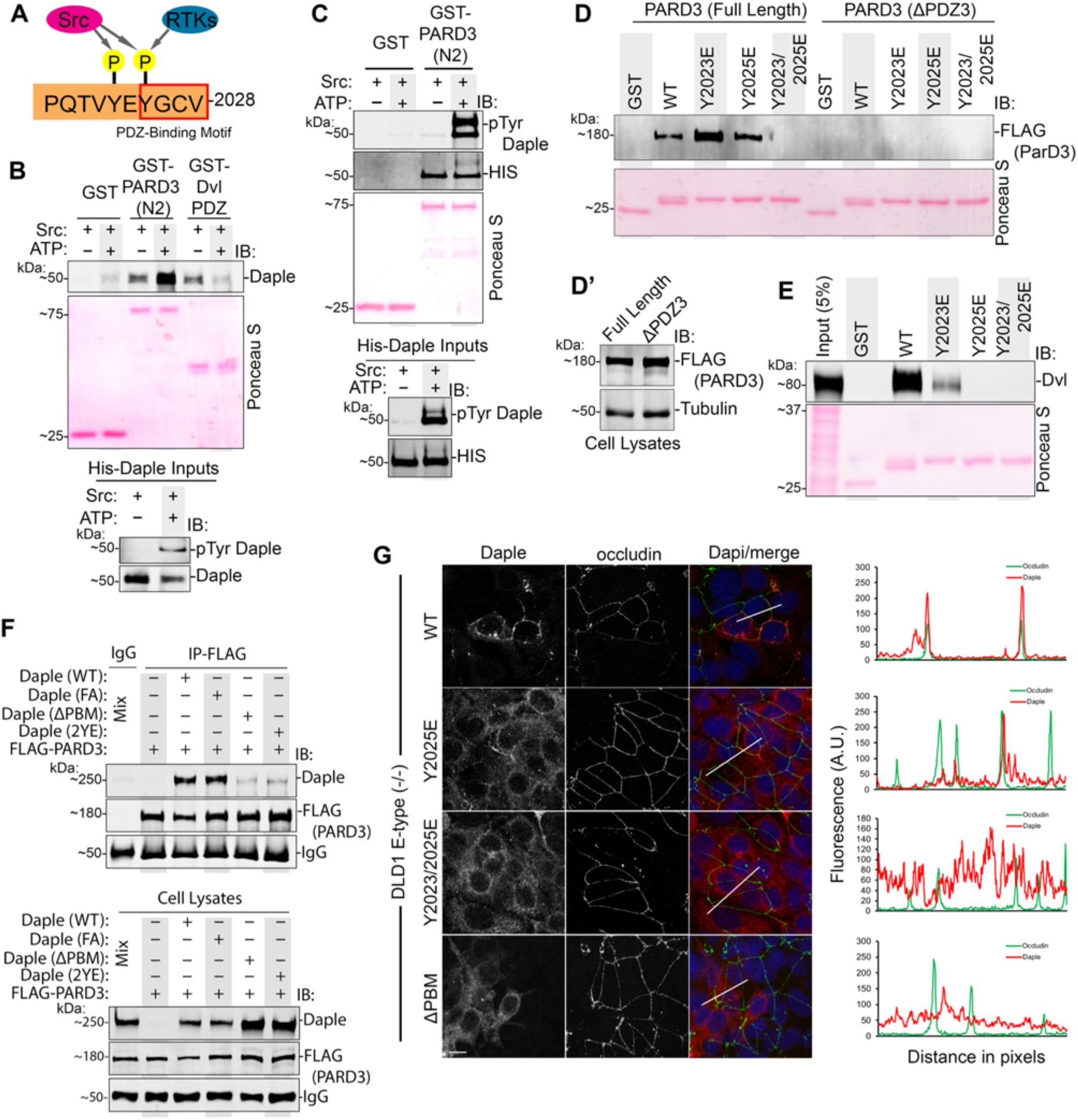
Tyrosine phosphorylation on Daple’s PBM regulates its ability to bind PARD3. **A)** Schematic summarizing published work of Daple’s PBM and its tyrosine phosphorylation site by RTKs (i.e. EGFR) or non-RTKs (i.e. Src) [10]. **B-C)** Daple-CT phosphorylated in vitro by recombinant Src kinase was used in GST-pulldown assays with PARD3-PDZ domains or Dvl-PDZ domain. Bound proteins are analyzed for Daple-CT using total (B) or phospho-specific (C) Daple antibody. **D-D’)** GST-tagged phosphomimicking Daple mutants (Y2023E, Y2025E, or double Y2023/2025E) were used in pulldown assays with lysates of transfected HEK293T as source for FLAG-PARD3-full length or PARD3-ΔPDZ3. Bound proteins are analyzed by immunoblotting in D. Input of cell lysates used in pulldown assay were analyzed by immunoblotting in D’. **E)** GST-tagged construct as in D, used in pulldown assays with lysates of transfected HEK293T as source for Dvl. Bound proteins are analyzed by immunoblotting. **F)** Co-immunoprecipitation assays investigating the binding between FLAG-PARD3 and various Daple mutants (WT, F1675A [FA], ΔPBM, or Y2023/2025E [2YE]). Proteins were exogenously expressed in HEK293T cells followed by cell lysis and IP using anti-FLAG antibody or control IgG. **G)** E-type (−/−) cells transiently expressing Daple-WT, Daple-Y2025E, or Daple-Y2023/2025E stated for Daple (*red*) and occludin (*green*). *Right*: RGB plot of indicated region indicated in merge panels.

Based on these findings, a picture emerges in which single phosphorylation at Y2023 or Y2025 may be sufficient to disrupt Daple•Dvl, but not Daple•PARD3 interaction, while dual site phosphorylation disrupts both. We previously showed that neither phosphoevent, alone or in combination, impact Daple’s ability to bind Gαi or FDZ [10] (**Figure 7A**). We conclude that graded phosphorylation of Daple’s PBM may regulate the composition of Daple-bound complexes, and may contribute, in part, to the dynamic spatial temporal control over their assembly/disassembly by tyrosine-based signals. To confirm if these events regulate planar migration, we generated “rescue” cell lines in which Daple-WT or two specific mutants that cannot bind PARD3 were stably expressed in Daple-depleted DLD1 cells. We noted that the levels of expression of Daple constructs were well above endogenous levels (**Figure S6E**) and that the constructs were expressed heterogeneously (**Figure S6F**). When we set out to carry out the polarized radial migration assays from the edge of tumor spheroids, we noted that all ‘rescue’ cell lines failed to form true compact spheroids like the parental line (**Figure S6G**). Because compact spheroids are a pre-requisite to study planar migration in the setting of intact junctions [26], and because of the importance of physiologic stoichiometry of signaling and scaffolding proteins [21, 38, 39], we did not proceed with radial migration assays in these lines. Future studies with sophisticated methods will be needed, e.g., genetic knock-in of the specific mutations so that endogenous levels can be achieved through its natural promoter. Given these limitations observed in cell lines, and in light of recently published *in vivo* findings using *Xenopus laevis [16]*, we conclude that the impact of phosphoevents we observe here on Daple-bound complexes *in vitro* may contribute to dynamic regulation of Daple-dependent signaling at cell–cell junctions in the epithelium.

**Figure 7.**
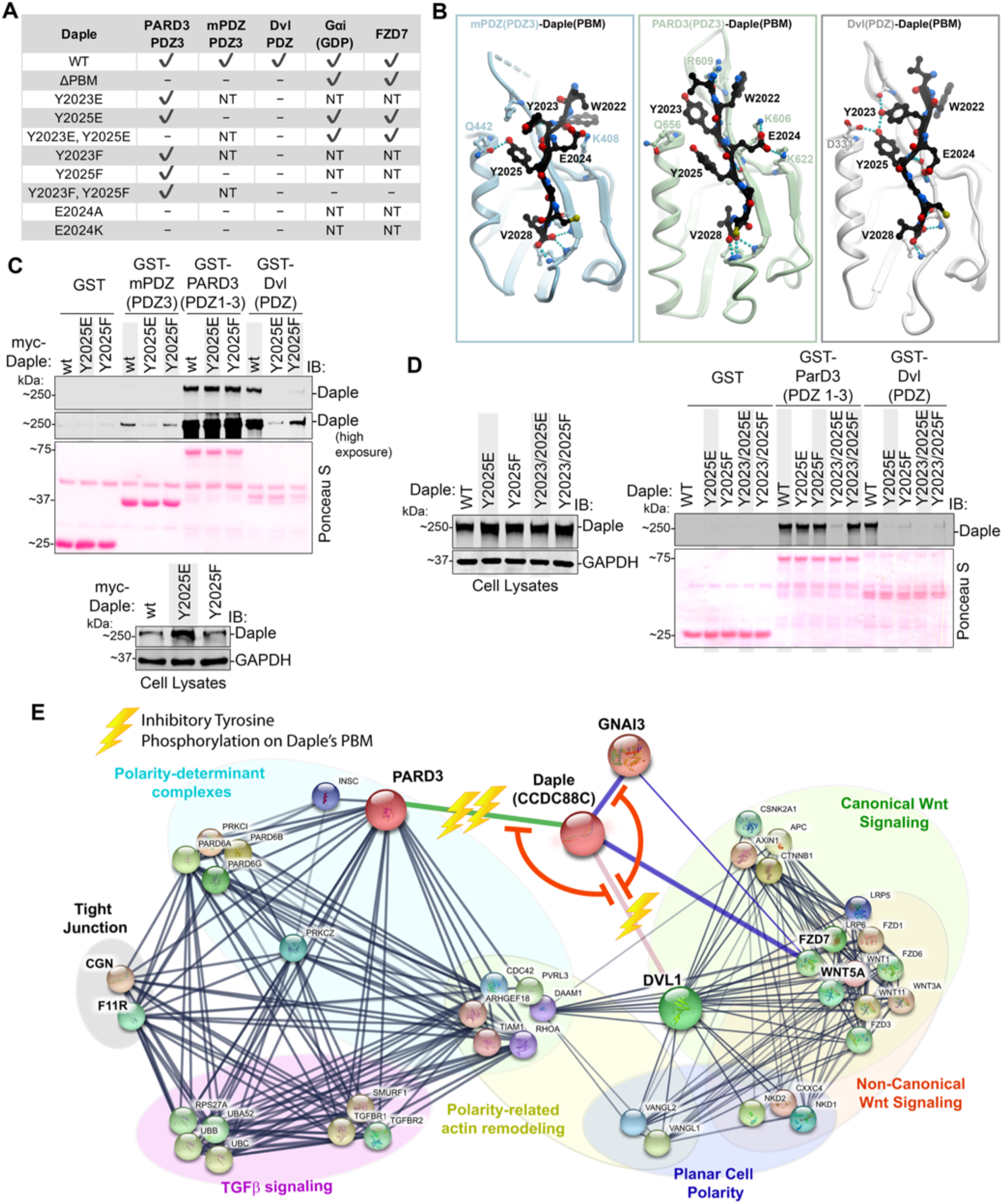
Summary of Findings: Phosphoregulated assembly of PARD3•Gαi3•Daple, Dvl•Daple, and Gαi3•Daple complexes at cell-cell junctions fine tunes planar cell migration. **A)** Table summarizing the interactome of Daple’s PBM with various PDZ domains, and the impact of various mutations tested in this work. NT, not tested. **B)** Homology models of Daple’s PBM bound to PDZ domains of PARD3, mPDZ, or Dvl. **C)** Pulldown assay carried out using GST-tagged PDZ domains of mPDZ (PDZ3), PARD3, and Dvl immobilized on glutathione S transferase beads and lysates of transiently transfected HEK293T cells as source for WT and mutants of myc-Daple (Daple-WT, phosphomimicking Daple-Y2025E, or unphosphorylated Daple-Y2025F). Immunoblots show bound proteins and confirm expression of Daple in cell lysates (input). **D)** GST pulldown assay carried out using GST-tagged PDZ domains of PARD3 and Dvl and lysates of transiently transfected HEK293T cells as source for WT and mutants of myc-Daple. Bound proteins were visualized by immunoblotting. **E)** A protein-protein interaction network built using STRING [40, 41] depicts how Daple may regulate various closely related biological processes via its interactions with PARD3, Dvl, and Gαi. Mutual exclusivity between complexes (red lines) and the impact of tyrosine phosphorylation in each interaction (bolt, inhibitory effect) is shown.

### Homology modeling reveals subtle differences between various Daple-PBM•PDZ complexes

To rationalize why some Daple-PBM•PDZ interactions were more sensitive to tyrosine phosphorylations (Dvl and mPDZ) than others (PARD3), we carried out 3D homology-modeling using a previously solved structure of the PDZ domains of PARD3, Dvl and mPDZ, in complex with peptide ligands. Both Y2023 and Y2025 form hydrogen bonds (H-bonds) with Dvl, which explains why the removal of the hydroxyl group (through phosphorylation, Y>E, or Y>F mutation) on either tyrosine abolishes Daple interaction with Dvl (**Figure 7B**). Only Y2025 on Daple forms a H-bond with mPDZ (to Q442), explaining similar detrimental effects of Y2025E and Y2025F on mPDZ binding to Daple. Similar to what was observed on Dvl, the Y2025E mutant showed a greater decrease in binding compared to Y2025F (**Figure 7C**). Homology modeling studies suggested that neither of the tyrosine residues contributes substantially to Daple’s interaction with the third PDZ domain of PARD3. This explains why mutation of either Y2023 or Y2025 to E or to F does not cause a loss of binding between Daple and PARD3 (**Figure 7D**). The increase in binding between PARD3 and Daple-Y2023E mutant and phosphorylated Daple can be explained by the increase in charge interaction between Y2023 and R609 on PARD3 (**Figure 6B-D** and **Figure 7B**).

The models also shed additional insights into the subtle differences in Daple’s ability to bind diverse PDZ domains: instead of the tyrosines, the key determinant of Daple binding to PARD3 and mPDZ is E2024 which forms a salt bridge with K606 and K622 (PARD3) and with K408 (mPDZ). Because Dvl lacks basic residues in the corresponding positions, the salt bridge is absent in the Daple•Dvl complex; however, E2024 appears to engage with the backbone of I282 (Dvl) via H-bonding. In keeping with these modeling-based predictions, GST-Daple-PBM with a E2024 mutation to Alanine (A) or Lysine (K) also abolished binding to PARD3 and Dvl (**Figure S7A-B**). In the reciprocal assays, when GST-PDZ protein was used to pulldown full length Daple form cell lysates, Daple-E2024A and E2024K mutants showed decreased binding to all three PDZ proteins, albeit to variable degrees (**Figure S7C**). The varying degrees of Daple•PARD3 or Daple•Dvl interactions observed in pulldown assays interchangeably using one binding partner as bacterially expressed GST protein and another from cell lysates suggest that the interaction in cells maybe more complex (i.e. subjected to additional posttranslational modifications) than what can be reconstituted in *in vitro* studies.

Together, our findings indicate that despite the overall shared modality of binding between Daple and the three PDZ proteins, subtle differences exist between the Daple-PBM•PDZ interface assembled in each case. These differences could account for their differential sensitivity to disruption by tyrosine phosphorylations at the two sites on Daple’s PBM. Furthermore, a STRING [40, 41] protein-protein interaction network shows that the disruptive phosphoevents on Daple’s PBM could provide a molecular basis for how tyrosine-based signals can initiate or terminate conditional and contextual scaffolding of key proteins within discrete pathways and cellular processes (**Figure 7E**).

## Conclusion

The major discovery we report here is that junction-localized Daple scaffolds PDZ-containing polarity regulator PARD3 to maintain contact-triggered polarization during migration in 2D planes. We also demonstrate how growth factors may shape these scaffolding functions of Daple via their ability to trigger two key tyrosine phosphoevents targeting Daple’s PBM. These phosphoevents dictate the localization of Daple, as well as its interactome: Daple that is not phosphorylated or hypophosphorylated (i.e., only on one tyrosine) may bind PARD3 and localize to the junctions, however, Daple that is hyperphosphorylated (i.e., on both tyrosines) may not bind either; instead, it localizes to the cytoplasm and may continue to modulate G protein signaling downstream of Wnt/FZD. Overall, our findings show that unphosphorylated Daple has maximal localization at cell-cell junctions; dually-phosphorylated Daple (by Src family of kinases) cannot localize to junctions; the mono-phosphorylated Daple has an intermediate hypomorphic characteristic.

These mechanistic studies, in conjunction with our prior work elucidating the effects of growth factor RTKs and growth factor stimulated non-RTKs on Daple [10], support the following working model in which the Daple•PARD3 interaction may serve as a critical determinant of localization of Daple at cell junctions. Once at junctions, Daple may orchestrate what appears to be a graded response to varying concentrations of growth factors: When growth factor concentrations are low (presumably triggering low-grade RTK activation), hypophosphorylated Daple bound to PARD3 promotes contact-triggered polarization during migration, however, when growth factors are high, hyperphosphorylated Daple dissociates from PARD3 and potentiates contact-free cell scattering.

Our findings also shed light into the two seemingly opposite ways in which Daple influences cancer initiation and progression; it acts as tumor suppressor in the normal epithelium and during early stages and, while supporting EMT/invasion during late stages [9, 42]. This bi-faceted role is shared by two other prominent signaling pathways—TGFβ and the non-canonical β-catenin-independent signaling that is triggered by Wnt5A/FZD7 [43, 44]. By scaffolding both these pathways to cell junctions (see **Figure 7E**), Daple may enable these growth factor signals to support seemingly opposite responses in cells with or without stable junctions. Cells with junctions, such as normal epithelium or in early-staged well-differentiated tumors will elicit contact-triggered orientation and contact-dependent planar migration. By contrast, cells without junctions, such as poorly-differentiated tumors, will elicit contact-free migration/EMT. Thus, it is possible that the seemingly opposing roles of Daple may be due to the presence or absence of cell-cell junctions and Daple’s localization to those junctions.

As for the relevance of our findings during development, it is noteworthy that mutations in Daple (that eliminate its PBM) have been identified in patients with non-syndromic hydrocephalus and depletion of Daple in animal models lead to hydrocephalus as well as ear/hearing defects [14–17, 45]. Underlying both conditions is a failure to polarize the cells and position the cilia. Our work provides mechanistic insights into how Daple may maintain contact-triggered cellular orientation in polarized cells with intact junctions. It is possible that the mechanisms we delineate here in colon epithelial cells are fundamental and hold true in other instances.

## Acknowledgements

This paper was supported by NIH CA238042, CA100768 and CA160911 (to P.G). J.E was supported by an NCI/NIH-funded Cancer Biology, Informatics & Omics (CBIO) Training Program (T32 CA067754) and a Postdoctoral Fellowship from the American Cancer Society (PF-18-101-01-CSM). I.K. was supported by NIH grants GM071872, AI118985, and GM117424. We thank Lee Swanson, Ying Dunkel, Nina Sun, Navin Rajapakse, Julie Choi, and Camille Delbrook for technical support in this work. We also thank Dr. Soumita Das at the UC San Diego HUMANOID CoRE for access to human organoids.

## Author Contributions

J.E, A.S, and P.G designed, performed and analyzed most of the experiments in this work. I.K generated the homology model for Daple-bound PARD3, mPDZ, and Dvl, and provided structure-based guidance to study Daple. M.G performed the mass spectrometry analysis. J.E and P.G conceived the project and wrote the manuscript.

## Declaration of Interests

The authors declare no competing interests.

