## Supplemental Material for "Tyrosine-based Signals Regulate the assembly of Daple•PARD3 Complex at Cell-cell Junctions during Polarized Planar Cell Migration"

#### INVENTORY OF SUPPLEMENTAL INFORMATION

**Supplemental Information includes:**

- STAR *Methods (Transparent Methods)
- Supplementary figures with legends. **S1 to S7**
- References Cited

**STAR *Methods**

- KEY RESOURCE TABLE
- CONTACT FOR REAGENNT AND RESOURCE SHARING
- EXPERIMENTAL MODEL AND SUBJECT DETAILS
    - Human IHC Samples
    - Cell Lines (DLD1, CaCo-2, SW480, HEK293T, HeLa, MDCK)
    - Human Enteroids
- METHOD DETAILS
    - Cell culture and DLD1 E-type and R-type isolation
    - Isolation, expansion and culture of organoids from human colons
    - Immunofluorescence and Confocal Microscopy, Image analysis
    - Image Processing
    - Cell Fractionation
    - Immunoblotting
    - Daple CRISPR/Cas9 Gene Editing and Validation
    - Transwell Migration Assay
    - Anchorage-dependent and Anchorage-independent Colony Growth Assays
    - Measurement of trans-epithelial electrical resistance (TEER)
    - Biotin Proximity Labeling
    - In Gel Digest
    - LC-MS analysis
    - Gene Ontology analysis
    - Recombinant Protein Purification
    - In Vitro GST-Pulldown and In-cellulo Co-immunoprecipitation (CoIP) Assays
    - In Vitro Kinase Assay
    - Spheroid formation and radial migration assay on Fibronectin
    - Molecular modeling
- QUANTIFICATION AND STATISTICAL ANALYSIS
    - Statistical Analysis
    - Replications
- DATA AND SOFTWARE AVAILABILITY

#### Key Resource Table

| REAGENT or RESOURCE | SOURCE | IDENTIFIER |
|---|---|---|
| **Antibodies** | | |
| Rabbit polyclonal anti-Daple | Millipore Sigma | ABS515 |
| Rabbit polyclonal anti-phospho Daple | 21st Century Biochemicals | N/A |
| Rabbit polyclonal anti-Gαi3 (C-10) | Santa Cruz Biotechnology | N/A |
| Rabbit polyclonal anti-ZO1 | GeneTex | GTX108613 |
| Rabbit polyclonal anti-ZO1 | GeneTex | GTX108627 |
| Rabbit polyclonal anti-PARD3 | Proteintech | 11085-1-AP |
| Rabbit polyclonal anti-E-cadheren | Santa Cruz Biotechnology | sc-7870 |
| Rabbit polyclonal anti-β-tubulin | Santa Cruz Biotechnology | sc-9104 |
| Rabbit polyclonal anti-GM130 | Cell Signaling Technology | 12480S |
| Mouse Monoclonal anti-occludin | ThermoFisher Scientific | 33-1500 |
| Mouse monoclonal anti-β-catenin | Santa Cruz Biotechnology | sc-7963 |
| Mouse monoclonal anti-GAPDH | Santa Cruz Biotechnology | sc-365062 |

| | | |
|---|---|---|
| Mouse monoclonal anti-αE-catenin | Santa Cruz Biotechnology | sc-9988 |
| Mouse monoclonal anti-FLAG | Millipore Sigma | MAB3118 |
| Mouse monoclonal anti-FLAG (hybridoma) | Purified in house | N/A |
| Mouse monoclonal anti-HIS | GenScript | A00186-100 |
| Mouse monoclonal anti-GST | GenScript | A00865 |
| Mouse monoclonal anti-Myc | Cell Signaling Technology | 2276S |
| Mouse monoclonal anti-Myc (hybridoma) | Purified in house | N/A |
| Mouse monoclonal anti-phosphotyrosine | BD Biosciences | 610000 |
| Mouse monoclonal anti-Dvl | Santa Cruz Biotechnology | sc-166303 |
| Mouse monoclonal anti-PARD3 | Novus Biologicals | MAB8030 |
| Mouse monoclonal anti-α-tubulin | Santa Cruz Biotechnology | sc-5286 |
| Goat anti-Rabbit IgG (680) | LI-COR Biosciences | 926-68071 |
| Goat anti-Rabbit IgG, Alexa Fluor 594 conjugated | ThermoFisher Scientific | A11072 |
| Goat anti-Mouse IgG (800) | LI-COR Biosciences | 926-32210 |
| Goat anti-Mouse IgG, Alexa Fluor 488 conjugated | ThermoFisher Scientific | A11017 |
| **Biological Samples and Cell Lines** | | |
| DLD1 | ATCC | CCL-221 |
| CaCo-2 | ATCC | HTB-37 |
| Sw480 | ATCC | CCL-228 |
| MDCK | ATCC | CCL-34 |
| HeLa | ATCC | CCL-2 |
| HEK293T | ATCC | CRL-11268 |
| Colonoids (healthy normal) | HUMANOID (Human organoid research Core, UCSD) | n/a |
| **Chemicals, Recombinant Proteins, and Plasmids** | | |
| Streptavidin, Alexa Fluor® 680 conjugate | ThermoFisher Scientific | S21378 |
| Streptavidin, Alexa Fluor® 594 conjugate | ThermoFisher Scientific | S11227 |
| Streptavidin Magnetic Beads | ThermoFisher Scientific | 88816 |
| Biotin | Sigma-Aldrich | B4639-500MG |
| DAPI (4',6-Diamidino-2-Phenylindole, Dilactate) | Thermo Fisher Scientific | D3571 |
| Fibronectin | Thermo Fisher Scientific | PHE0023 |
| TPA (12-O-Tetradecanoylphorbol 13-acetate, PMA, Phorbol 12-myristate 13-acetate) | Sigma-Aldrich | P1585 |
| pSpCas9(BB)-2A-Puro (PX459) V2.0 | Addgene | 62988 |
| pME-BirA | *This paper* | N/A |
| p3E-Daple | *This paper* | N/A |
| p3E-Daple-CT-WT (a.a. 1650-2028) | *This paper* | N/A |
| p3E-Daple-CT-ΔPBM (a.a. 1650-2025) | *This paper* | N/A |
| pcsDest2-BirA-Daple | *This paper* | N/A |
| pDEST17-Daple- CT-WT (a.a. 1650-2028) | *This paper* | N/A |

| | | |
|---|---|---|
| pDEST17-Daple-CT-ΔPBM (a.a. 1650-2025) | *This paper* | N/A |
| pGEX-4T-Daple-CT-WT (a.a. 2000-2018) | *This paper* | N/A |
| pGEX-4T-Daple-CT-ΔPBM (a.a. 2000-2018) | *This paper* | N/A |
| pGEX-4T-Daple-CT-Y2023E (a.a. 2000-2018) | *This paper* | N/A |
| pGEX-4T-Daple-CT-Y2025E (a.a. 2000-2018) | *This paper* | N/A |
| pGEX-4T-Daple-CT-Y2023/2025E (a.a. 2000-2018) | *This paper* | N/A |
| pGEX-4T-Daple-CT-E2024A (a.a. 2000-2018) | *This paper* | N/A |
| pGEX-4T-Daple-CT-E2024K (a.a. 2000-2028) | *This paper* | N/A |
| pGEX-4T-Daple-CT-WT (a.a. 1650-2028) | Aznar, et. al., 2015 | N/A |
| pET28b-Daple-CT-WT (a.a. 1650-2028) | Aznar, et. al., 2015 | N/A |
| myc-pcDNA 3.1 (+) - Daple-WT (full length) | Aznar, et. al., 2015 | N/A |
| myc-pcDNA 3.1 (+) - Daple-ΔPBM (full length) | Aznar, et. al., 2015 | N/A |
| myc-pcDNA 3.1 (+) - Daple-FA (full length) | Aznar, et. al., 2015 | N/A |
| myc-pcDNA 3.1 (+) - Daple-Y2025E (full length) | Aznar, et. al., 2018 | N/A |
| myc-pcDNA 3.1 (+) - Daple-Y2025F (full length) | Aznar, et. al., 2018 | N/A |
| myc-pcDNA 3.1 (+) - Daple-2YE (full length) | Aznar, et. al., 2018 | N/A |
| myc-pcDNA 3.1 (+) - Daple-2YF (full length) | Aznar, et. al., 2018 | N/A |
| pGEX-5X-1 mPDZ PDZ 1-13 | Baliova M, et. al., 2014 | N/A |
| pcDNA3.1-N-term FLAG-PARD3 (full length) | Peng Zhang, et. al., 2016 | N/A |
| pcDNA3.1-N-term FLAG-PARD3 (ΔPDZ 1) | Peng Zhang, et. al., 2016 | N/A |
| pcDNA3.1-N-term FLAG-PARD3 (ΔPDZ 2) | Peng Zhang, et. al., 2016 | N/A |
| pcDNA3.1-N-term FLAG-PARD3 (ΔPDZ 3) | Peng Zhang, et. al., 2016 | N/A |
| GST-PARD3-1N | Norimichi Itoh, et. al., 2010 | N/A |
| GST-PARD3-2N | Norimichi Itoh, et. al., 2010 | N/A |
| GST-PARD3-3N | Norimichi Itoh, et. al., 2010 | N/A |
| GST-PARD3-4N | Norimichi Itoh, et. al., 2010 | N/A |
| GST-ZO-1 PDZ 1-3 | Kris Meerschaert, et. al., 2009 | N/A |
| GST-ZO-1 PDZ 1 | Kris Meerschaert, et. al., 2009 | N/A |
| GST-ZO-1 PDZ 2 | Kris Meerschaert, et. al., 2009 | N/A |
| GST-ZO-1 PDZ 3 | Kris Meerschaert, et. al., 2009 | N/A |
| GST-ZO-2 PDZ 1 | Kris Meerschaert, et. al., 2009 | N/A |
| GST-ZO-2 PDZ 2 | Kris Meerschaert, et. al., 2009 | N/A |
| GST-ZO-2 PDZ 3 | Kris Meerschaert, et. al., 2009 | N/A |
| pGEX4T3-GIPC-PDZ | Tal Varsano, et. al., 2012 | N/A |
| GST-mPDZ (PDZ 1 through 13) | Martina Baliova, et. al., 2014 | N/A |

| Software | | |
|---|---|---|
| ImageJ | National Institute of Health | https://imagej.net/Welcome |
| DAVID 6.8 | DAVID Bioinformatics Resources | https://david.ncifcrf.gov/home.jsp |
| String | Snel B, et. al., 2000 | string-db.org |
| OncoLnc | Anaya J. 2016 | http://www.oncolnc.org |
| Molsoft ICM v3.8-6 | Molsoft LLC | http://www.molsoft.com/index.html |

*Cell culture and DLD1 E-type and R-type isolation*

All DLD1 cells were cultured using RMPI media containing 10% FBS. Cells were routinely passaged at a dilution of 1:5 to 1:10. HEK293T cells were cultured using DMEM media containing 10% FBS and routinely passaged at a dilution of 1:10.

For E-type and R-type isolation, parental stock of DLD1 cells were resuspended into single cells and plated sparsely onto 10cm tissue culture plates so that individual colonies can be picked. E-type and R-type cells can be distinguished morphologically under microscope through analyzing the amount of light passing through between cells. Further validation of proper E-type and R-type isolation was confirmed through immunoblotting for $\alpha$-catenin.

*Isolation, expansion and culture of organoids from human colons*

Intestinal crypts were isolated from the colonic tissue specimen by digesting with Collagenase type I [2 mg/ml; Life Technologies Corporation, NY) and cultured in stem-cell enriched conditioned media with WNT 3a, R-spondin and Noggin [1-3]. Briefly, after digestion with Collagenase the crypts were filtered with a cell strainer and washed with DMEM/F12 with HEPES, supplemented with 10% FBS. After adding collagenase I solution containing gentamicin (50 $\mu$g/ml, Life Technologies Corporation, NY) and mixing thoroughly, the plate was incubated at 37° C inside a $CO_2$ incubator for 10 min with vigorous pipetting between incubations and monitoring constantly by light microscopy to confirm by direct observation the dislodgement of the intestinal crypts from the tissues. The collagenase was inactivated with DMEM/F12 with HEPES, supplemented with 10% FBS and filtered using a 70 $\mu$m cell strainer over a 50 ml centrifuge tube. Filtered tissue was spun down at 200xg for 5 min, and the media was aspirated. The epithelial units were suspended in matrigel. Cell-matrigel suspension (15 $\mu$l) was placed at the center of the 24-well plate on ice and placed on the incubator upside-down for polymerization. After 10 min, 500 µl of 50% conditioned media (CM) was added. CM was prepared from L-WRN cells [ATCC® CRL-3276™ [3]] with Wnt3a, R-spondin and Noggin. Y27632 (ROCK inhibitor, 10 µM) and SB431542 (an inhibitor for TGF-β type I receptor, 10 µM) were added to the media. For human enteroids, additional supplements (purchased from Cell Applications Inc. San Diego, CA) were added to the above media. The medium was changed every 2-3 days and the enteroids were expanded as needed for experimentation.

*Immunofluorescence and Confocal Microscopy, Image analysis*

Cells or organoids were fixed at using -20°C methanol for 20 to 30 mins, rinse with PBS, then permeabilized for 1hr using blocking/permeabilization buffer (0.4% Triton X-100 and 2 mg/ml BSA dissolved in PBS). Primary antibody and secondary antibody were diluted in blocking buffer and incubation was carried out for 1 hr each. Coverslips were mounted using Prolong Gold (invitrogen) and imaged using a Leica SPE CTR4000 with a 63X objected and 1AU aperture. Where z-stacks was performed, images were taken at 0.5 μm slices.

*Image Processing*

All images were processed on ImageJ software (NIH) and assembled into figure panels using Photoshop and Illustrator (Adobe). RGB plots were generated by exporting ROI plot values and graphing into Excel (Microsoft).

*Cell Fractionation*

Cells expressing were harvested and suspended in homogenization buffer (10 mM sodium phosphate buffer [pH 7.2], 1 mM $MgCl_2$, 30 mM NaCl, 1 mM DTT, and 0.5 mM phenylmethylsulfonyl fluoride, supplemented with protease and phosphatase inhibitors), and homogenized using a 30-gauge needle. Crude membranes from the homogenate were pelleted by centrifugation of post-nuclear supernatant at 100,000 × *g* for 60 min at 4°C in a TLA-41 fixed-angle rotor in a TLA-100 table-top ultracentrifuge (Beckman Coulter, Krefeld, Germany). Pelleted membranes were washed in homogenization buffer before resuspension in cell lysis buffer containing 0.4% Tx-100.

*Immunoblotting*

For immunoblotting, protein samples were prepared in Laemmli sample buffer, separated by SDS-PAGE and transferred onto 0.4μm PVDF membrane (Millipore). After transfer, membranes were blocked with 5% Non-fat milk or 5% BSA (for phosphoprotein blotting) in PBS. Primary antibodies were prepared in blocking buffer contain 0.1% Tween-20 and incubated with blots overnight at 4°C. After primary antibody incubation, blots were incubated with secondary antibodies for one hour at room temperature and imaged using a Li-Cor Odyssey imaging system.

*Daple CRISPR/Cas9 Gene Editing and Validation*

Daple target genomic DNA sequence was cloned into PX-459 vector and transfected into DLD1 cells. Puromycin was added to cells 30 hours post transfection for selection. When untransfected control plates showed 95 to 100% cell death, cells were washed with PBS and fresh media (with no puromycin) was added to allow cells to recover for 8 hours. Following recovery, cells were resuspended and plated sparsely onto 10 cm plates so that individual cell colonies could be picked into 12-well plates and screened for indels.

In order to identify cell clones harboring mutations in gene coding sequence, genomic DNA was extracted using 50 mM NaOH and boiling at 95°C for 60mins. After extraction, pH was neutralized by the addition of 1.0 M Tris-pH 8.0 (10% volume). Crude genomic extract was then used in PCR reactions with primers flanking the protospacer adjacent motif (PAM) sequence. Amplicons were analyzed for indels using TBE-PAGE gels. Sequence of mutation was determined by cloning amplicons into sequencing vector using TOPO-TA cloning (Invitrogen).

*Transwell Migration Assay*

Transwell plates (24-well; 8.0 μm pore size; Corning) were used for chemotactic cell migration assays. Cells were trypsinized, resuspended in media containing no FBS, and placed into Transwells (200,000 cells/well). Transwells were then placed into plates containing no FBS and cells were allowed to settle for 30 mins. Inserts were then transferred into chambers containing 2% FBS overnight to trigger chemotactic migration. For fixing and staining of cells, transwells were first rinsed with PBS, then fixed using 3% paraformaldehyde in PBS for 15 mins. After fixation, transwells was rinsed with PBS, then permeabilized using 100% methanol for 15 mins, followed by rinsing in distilled water. Staining of cells were carried about by submerging transwell membrane in a 2% crystal violet solution for 30 mins, then rinse using distilled water. Non-migrated cells in upper chamber was removed using cotton swab.

*Anchorage-dependent and Anchorage-independent Colony Growth Assays*

Anchorage-dependent growth assays were monitored on regular tissue culture plastics. Cells were resuspended and plated at 1,000 cells per well in a 6-well plate and incubated for approximately 10 days in 10% FBS media. Cells were then fix and permeabilized using 100% methanol. Cells were stained with 2% crystal violet and then rinsed using distilled water.

Anchorage-independent growth was performed by growing cells in a suspension of agar. A base layer of agar was prepared by dissolving 0.6% agarose in RPMI media containing 10% FBS, and then placing 3.0 ml of the solution into a 6.0 cm plate. Base layer was allowed to cool and set at room temperature for 1 hr before cell layer was added. For the cell layer, 5,000 cells were resuspended in 3.0 ml of 0.5% low melting agarose dissolved in RPMI media containing 10% FBS and then slowly placed on top of base layer with the precaution of avoiding air bubbles. Cell layer was allowed to cool and set at room temperature for 60 mins prior to transferring to 4°C for 10 mins. Plates were then placed into 37°C incubator with 5% $CO_2$ overnight, then 0.5 ml of RPMI media with 10% FBS was added on top and occasionally replaced to keep cells hydrated. Cells were grown for approximately 3 to 4 weeks. Cells were stained using 0.05% crystal violet dissolved in 10% ethanol. Staining solution was placed on cells for 60 mins at room temperature. Solution was then gently removed, and cells were washed several times with distilled water.

For both aforementioned colony growth assays, images were acquired by light microscopy and colonies were counted using ImageJ (NIH).

*Measurement of trans-epithelial electrical resistance (TEER)*

TEER of DLD1 cells was measured by culturing cells on a 0.4 μm pore size 12-mm polycarbonate Transwell Filter (Corning) followed by measurements using an epithelial voltohmmeter (Millicel-ERS resistance meter, Millipore). Cells were plated at approximately 1.5 X $10^5$ cells per filter. Cells were grown for 5 days to allow for the establishment of a confluent monolayer.

*Biotin Proximity Labeling*

Cells were plated 24 hrs prior to transfection with Daple-BirA construct. Thirty hours post transfection, cells were incubated with 50 μM biotin for 16 hrs. Prior to lysis, cells were rinsed two times with PBS, then lysed by resuspending in lysis buffer containing (50 mM Tris, pH 7.4, 500 mM NaCl, 0.4% SDS, 1 mM dithiothreitol, 2% Triton X-100, and 1× Complete protease inhibitor) and sonication. Cell lysates were then cleared through centrifugation at 20,000 X g for 20 mins. Supernatant was then collected and incubated with Streptavidin Dynabeads overnight at 4°C. Beads were then washed twice with 2% SDS, once with wash buffer 1 (0.1% deoxycholate, 1% Triton X-100, 500 mM NaCl, 1 mM EDTA, and 50 mM HEPES, pH 7.5), followed with once wash using wash buffer 2 (250 mM LiCl, 0.5% NP-40, 0.5% deoxycholate, 1 mM EDTA, and 10 mM Tris, pH 8.0), and once with 50mM Tris pH 8.0. Biotinylated complexes were then eluted using sample buffer containing excess biotin and heating to 100°C.

For co-immunoprecipitation of protein-protein complexes from cell lysates, cells were first lysed in cell lysis buffer (20 mM HEPES, pH 7.2, 5 mM Mg-acetate, 125 mM K-acetate, 0.4% Triton X-100, 1 mM DTT, 500 µM sodium orthovanadate, phosphatase inhibitor cocktail (Sigma-Aldrich) and protease inhibitor cocktail (Roche Life Science)) using a 28G syringe, followed by centrifugation at 10,000Xg for 10mins. Cleared supernatant was then used in binding reaction with immobilized GST-proteins for 4 hours at 4°C. After binding, bound complexes were washed four times with 1 ml phosphate wash buffer (4.3 mM Na2HPO4, 1.4 mM KH2PO4, pH 7.4, 137 mM NaCl, 2.7 mM KCl, 0.1% (v:v) Tween 20, 10 mM MgCl2, 5 mM EDTA, 2 mM DTT, 0.5 mM sodium orthovanadate). Bound proteins were then eluted through boiling at 100°C in sample buffer. Prior to mass spectrometry identification, eluted samples were run on SDS-PAGE and proteins were extracted by in gel digest.

*In Gel Digest*

Protein digest and mass spectrometry was perform as previously described[4]. The gel slices were cut to 1mm by 1 mm cubes and destained 3 times by first washing with 100 ul of 100 mM ammonium bicarbonate for 15 minutes, followed by addition of the same volume of acetonitrile (ACN) for 15 minutes. The supernatant was and samples were dried in a speedvac. Samples were then reduced by mixing with 200 µl of 100 mM ammonium bicarbonate-10 mM DTT and incubated at 56C for 30 minutes. The liquid was removed and 200 ul of 100 mM ammonium bicarbonate-55mM iodoacetamide was added to gel pieces and incubated at room temperature in the dark for 20 minutes. After the removal of the supernatant and one wash with 100 mM ammonium bicarbonate for 15 minutes, same volume of ACN was added to dehydrate the gel pieces. The solution was then removed and samples were dried in a speedvac. For digestion, enough solution of ice-cold trypsin (0.01 ug/ul) in 50 mM ammonium bicarbonate was added to cover the gel pieces and set on ice for 30 min. After complete rehydration, the excess trypsin solution was removed, replaced with fresh 50 mM ammonium bicarbonate, and left overnight at 37°C. The peptides were extracted twice by the addition of 50 µl of 0.2% formic acid and 5 % ACN and vortex mixing at room temperature for 30 min. The supernatant was removed and saved. A total of 50 µl of 50% ACN-0.2% formic acid was added to the sample, which was vortexed again at room temperature for 30 min. The supernatant was removed and combined with the supernatant from the first extraction. The combined extractions are analyzed directly by liquid chromatography (LC) in combination with tandem mass spectroscopy (MS/MS) using electrospray ionization.

*LC-MS analysis*

Trypsin-digested peptides were analyzed by ultra high pressure liquid chromatography (UPLC) coupled with tandem mass spectroscopy (LC-MS/MS) using nano-spray ionization. The nanospray ionization experiments were performed using a Orbitrap fusion Lumos hybrid mass spectrometer (Thermo) interfaced with nano-scale reversed-phase UPLC (Thermo Dionex UltiMate™ 3000 RSLC nano System) using a 25 cm, 75-micron ID glass capillary packed with 1.7-µm C18 (130) BEH™ beads (Waters corporation). Peptides were eluted from the C18 column into the mass spectrometer using a linear gradient (5–80%) of ACN (Acetonitrile) at a flow rate of 375 µl/min for 1h. The buffers used to create the ACN gradient were: Buffer A (98% $H_2O$, 2% ACN, 0.1% formic acid) and Buffer B (100% ACN, 0.1% formic acid). Mass spectrometer parameters are as follows; an MS1 survey scan using the orbitrap detector (mass range (m/z): 400-1500 (using quadrupole isolation), 120000 resolution setting, spray voltage of 2200 V, Ion transfer tube temperature of 275 C, AGC target of 400000, and maximum injection time of 50 ms) was followed by data dependent scans (top speed for most intense ions, with charge state set to only include +2-5 ions, and 5 second exclusion time, while selecting ions with minimal intensities of 50000 at in which the collision event was carried out in the high energy collision cell (HCD Collision Energy of 30%), and the fragment masses where analyzed in the ion trap mass analyzer (With ion trap scan rate of turbo, first mass m/z was 100, AGC Target 5000 and maximum injection time of 35ms). Protein identification and label free quantification was carried out using Peaks Studio 8.5 (Bioinformatics solutions Inc.)

*Gene Ontology Analysis*

Identified proteins unique to plus biotin samples, but not in minus biotin samples, were analyzed using DAVID and functional annotation was grouped by INTERPRO protein domains for GO analysis. Classification with p-value less than 0.5 was set as significant.

*Recombinant Protein Purification*

Both GST and His-tagged proteins were expressed in E. coli stain BL21 (DE3) and purified as previously described. Briefly, cultures were induced using 1mM IPTG overnight at 25°C. Cells were then pelleted and resuspended in either GST lysis buffer (25 mM Tris-HCl, pH 7.5, 20 mM NaCl, 1 mM EDTA, 20% (vol/vol) glycerol, 1% (vol/vol) Triton X-100, 2×protease inhibitor cocktail) or His lysis buffer (50 mM $NaH_2PO_4$ (pH 7.4), 300 mM NaCl, 10 mM imidazole, 1% (vol/vol) Triton X-100, 2×protease inhibitor cocktail). Cells were lysed by sonication, and lysates were cleared by centrifugation at 12,000 X g at 4°C for 30 mins. Supernatant was then affinity purified using glutathione-Sepharose 4B beads (GE Healthcare) or HisPur Cobalt Resin (Thermo Fisher Scientific), followed by elution, overnight dialysis in PBS, and then storage at -80°C.

*In Vitro GST-Pulldown and In-cellulo Co-immunoprecipitation (CoIP) Assays*

Purified GST-tagged proteins from E. coli were immobilized onto glutathione-Sepharose beads and incubated with binding buffer (50 mM Tris-HCl (pH 7.4), 100 mM NaCl, 0.4% (v:v) Nonidet P-40, 10 mM MgCl2, 5 mM EDTA, 2 mM DTT, 1X Complete protease inhibitor) for 60mins at room temperature. For GST-pulldown assays with recombinant proteins, the proteins were diluted in binding buffer and incubated with immobilized GST-proteins for 90mins at room temperature.

*In Vitro Kinase Assays*

In vitro kinase assays were performed using recombinant His-tagged Daple-CT (1650 to 2028) proteins purified from E. coli (BL21) and 50 ng of active GST-tagged recombinant Src Kinase (SignalChem, Canada). Substrates and kinase were mixed into tyrosine kinase buffer (60 mM HEPES pH 7.5, 5 mM MgCl2, 5 mM MnCl2, 3 μM sodium orthovanadate) and reaction was started by adding 1.0 mM ATP. Reactions was incubated at 25°C for 60 mins, and then used in subsequent pulldown assays or terminated using Laemmli Sample Buffer or boiling at 100°C.

*Spheroid Formation and Radial Migration Assays on Fibronectin*

Spheroids were produced by adding 4,000 cells into a 96-well plate precoated with 1% agarose and grown for 5 days, when well defined spheroids and necrotic core can be observed. For migration assay, the spheroids were transferred on coverslips coated with fibronectin (5 μg/ml) and the extent of radial migration was measured after 24 hrs by serial imaging of the spheroid by light microscopy. The area of migration was assessed by measuring surface area covered by cells and subtracting surface area of spheroids. Polarization towards the free-edge, and away from the tumor spheroid, was determine by the position of the Golgi towards the free-edge.

*Molecular modeling*

A structure of mouse PARD3 (PDZ3) with the C-terminus of mouse Cadherin-5 (QEELII, PDB 2koh [5]), and a structure of human mPDZ (PDZ3) with the C-terminus of the PM Ca-transporting ATPase 4 (ETSV, PDB 2iwn [6]) were used as docking templates. The Daple PBM peptide (2021-VWYEYGCV-2028) was modeled ab initio. Although many PDZ domains recognize their target PBM peptides via a canonical anti-parallel b-sheet interaction with alternating residue backbone

H-bonding pattern, this is not the case for Dvl (e.g. PDB 3cbx [7]). Consequently, the published model of Dvl2 in complex with Daple PBM [8] featured a kink in the peptide backbone. In this work, we explored the possibility of a similar kink in the complexes of Daple PBM with PARD3 (PDZ3) and mPDZ (PDZ3). For this, three options for H-bonding pattern between the backbones of the peptide and the target PDZ domain were tested by the modeling procedure: one skipping Daple V2028/hn bonding to Dvl I280/o (which corresponds to PARD3 V603/o and mPDZ I389/o), another skipping Daple Y2025/o and G2026/o bonding to Dvl I282/hn (which corresponds to PARD3 V605/hn and mPDZ I391/hn), and the third without H-bond skipping. For each H-bonding pattern, soft harmonic distance restraints were imposed between backbone atoms of Daple PBM peptide and the corresponding backbone atoms of the target PDZ domain. After this, the system consisting of the fully flexible peptide and the PDZ domain with flexible side-chains was thoroughly sampled using biased probability Monte Carlo sampling in internal coordinates as implemented in ICM[9]. The objective function included full-atom van der Waals term calculated using the Lennard-Jones potential and capped at 20 kcal/mol, hydrogen bonding term, electrostatics, torsional strain, and a penalty for the backbone distance restraints. Multiple poses generated by the docking procedure for each target domain were merged in a single list and rank-ordered by the predicted energy excluding the distance restraint penalty. Top-scoring conformation of each complex was selected for further analysis.

*Statistical Analysis and Replicates*

Graphs comparing E-type and R-type, with or without Daple, are represented as the mean ± standard error of the mean (SEM). Student's t-test was used to determine significance with P values of $< 0.05$ set as the threshold for statistical significance. Where statistical analysis was performed, experiments were performed in triplicates.

*Data and Software Availability*

We have not generated software and raw data of MS hits are available upon request from authors.

**Supplementary Figure Legends**

**Figure S1-Related to Figure 2. Daple localization and expression in colorectal cancer cell lines.**

**A)** CaCo-2 (*top*) and Sw480 (*bottom*) colorectal cancer cells were fixed with methanol, stained for Daple (*red*), occludin (*green*) and nucleus (DAPI; blue) and analyzed by confocal microscopy. Representative images are shown. Scale Bar, 25μm. **B)** A Kaplan Meier Plot was generated using OncoLnc (www.oncolnc.org) to compare the disease-free survival rates of patients with colorectal tumors with high vs. low levels of α-catenin expression. OncoLnc links The Cancer Genome Atlas (TCGA; NCI) survival data to mRNA levels [10]. High and low expression is defined as patients with top 25% and bottom 25% expression level, respectively. **C)** Parental DLD1 cells were diluted onto tissue culture plates to produce isolated colonies and imaged by light microscopy. Representative images of those colonies are shown. **D)** Whole cell lysates from clonally isolated colonies were analyzed for expression of Daple, E-cadherin, β-catenin, α-catenin, tubulin and GAPDH by immunoblotting (IB). **E)** EGTA (30 mins) was used to deplete calcium levels and disrupt cell junctions in DLD1 E-type cells. After treatment, cells were fixed and stained for Daple (*red*) and occludin (*green*) and assessed by confocal microscopy. Representative images are shown. Scale Bar, 25μm **F)** TPA (3 hours) was used to activate PKC and transiently restore junctions in R-type cells. Post treatment, cells were fixed and stained for Daple (*red*) and occludin (*green*). Although junctions were restored (as determined by occludin staining), Daple was not recruited to cell-cell junctions. Scale Bar, 25μm.

**Figure S2-Related to Figure 2. Subcellular fractionation studies on colorectal cancer cells reveal a pool of Daple that is on membrane and is detergent insoluble.**

**A)** Schematic and workflow for subcellular fractionation used in panels B (CaCo-2 and Sw480 cells) and C (DLD1 E-type and R-type cells). Nuclear free cell homogenates (PNS, post nuclear supernatant) in detergent free buffer was spun at 100,000 g to isolate cell membrane (P100, pellet 100,000 g) from cytosol (S100, supernatant 100,000 g). P100 fraction was then resuspended in buffer containing detergent, Triton Tx-100, and centrifuged at 120,000 g to isolate detergent soluble (Tx-100) from detergent insoluble (Insol.) fractions. **B-C)** Daple expression was determined in various cellular fractions by immunoblotting (IB). As controls, Gαi3 and E-cadherin was used to confirm membrane enrichment, whereas tubulin was used to determine cytosol enrichment. Membrane associated ZO-1 and PARD3 was observed in Tx-100 insoluble fractions whereas E-cadherin, β-catenin, and α-catenin was observed in both Tx-100 soluble and insoluble fractions. **C)** Lower panels show higher loading volume of P100, Tx-100, and Insoluble fractions of DLD1 E-type and R-type cells. Actin and Tubulin was used to confirm presence of cytoskeletal proteins. Caveolin-1 was used to confirm presence of lipid rafts.

**Figure S3-Related to Figure 2. Characterization and validation of Daple CRISPR/Cas9 knockout**

**A)** Schematic illustrating Cas9 target site on Exon 5 of Daple (*top*). PCR of region flanking target site using genomic DNA from various cell clones from Cas9 selection (*bottom*). **B)** Representative mutations identified through TOPO-TA cloning and sequencing of amplicons in (A). **C)** DLD1 E-type and R-type cell clones were fixed and stained for Daple (*red*). Scale bar, 25μm. **D)** Graph showing the volume of DLD1 E-type (+/+ and -/-) spheroids in panel E over time, as determined by ImageJ. **E)** Daple depleted DLD1 E-type and R-type cells were grown in agarose-coated plates for 7 days and serially imaged by light microscopy. Representative images of spheroids are shown. Scale bar, 1mm.

**Figure S4-Related to Figure 4. Validation of the Daple construct that was used in biotin proximity labeling.**

**A)** Schematic showing the domain composition of a myc-BirA-tagged Daple construct that was used for proximity labeling (Figure 5). **B)** HEK293T cells expressing myc-BirA-Daple were fixed and stained for Daple (*red*) or myc-tag (*green*) and analyzed by confocal microscopy. Representative images are shown. Scale bar, 25 μm. **C)** HEK293T cells were stained for endogenous Daple (red) and γ-tubulin (green). Scale bar, 5 μm. Asterisk = pericentriolar localization; arrow = PM localization. **D)** GST-pulldown assays were carried out using GST-tagged PDZ domain of Dvl and lysates of HEK293T cells with exogenously expressed myc-BirA-Daple. Immunoblots show myc-BirA-Daple bound to GST-Dvl-PDZ and confirm expression in cell lysates used in interaction assay. **E)** Co-Immunoprecipitation assays were carried out on lysates of HEK293T cells exogenously expressing myc-BirA-Daple and Dvl using anti-myc IgG or control IgG. Immunoblots show interaction of myc-BirA-Daple and Dvl in cells and confirm expression in cell lysates (input).

**Figure S5-Related to Figure 5. Daple binds directly to mPDZ; the interaction is mediated via Daple's PBM and the third PDZ module on mPDZ.**

**A)** Various GST-tagged PDZ domains on mPDZ were immobilized onto glutathione S transferase beads and used in an interaction assay with recombinant His-Daple-CT. Immunoblots show interaction of purified Daple-CT with the 3$^{rd}$, but not the other PDZ modules of mPDZ. **B)** mPDZ domains were bound to beads, as in (A), and used to pulldown myc-Daple from HEK293T cells. Immunoblots show bound proteins and confirm expression of Daple in cell lysates used in binding assay (input).

**Figure S6-Related to Figure 6. Impact of Daple and its tyrosine phosphorylation on binding and localization of PARD3.**

**A-B)** GST-tagged PDZ domain of PARD3 (A) or Dvl (B) were immobilized onto glutathione S transferase beads and used in an interaction assay with recombinant His-Daple-CT WT, Y2023E, Y2023F, Y2025E, or Y2025F. Interaction was determined through immunoblotting for Daple. **C)** GST-tagged PDZ domains of PARD3 used to pulldown ectopically expressed full length Daple-WT, Y2023E, Y2023F, Y2025E, Y2025F, Y2023/2025E, or Y2023/2025F in HEK293T cell lysates. Immunoblot for Daple show reveal loss of binding specifically in double phosphomimicking mutants. **D)** E-type (+/+) and E-type (-/-) cells were stained for Daple (*red*) and PARD3 (*green*). Scale Bar, 5μm. *Right:* RGB plot of indicated region in merge panels. **E)** Stable rescue lines of DLD1 E-type (-/-) cells expressing recombinant Daple-WT, ΔPBM, or Y2023/2025E were immunoblotted to determine expression levels. **F)** Stable lines, as described in F, were grown as spheroids as described in E and imaged on day 7. Selected regions show an inability to produce a compact spheroid and the presence of loose cells at the periphery. **G)** Stable rescue lines, as in F, were stained for Daple (red). Scale bar, 10μm.

**Figure S7-Related to Figure 7. Impact of E2024 on Daple's ability to bind PDZ-domain containing proteins.**

**A-B)** GST-tagged Daple-PBM-WT, E2024A, and E2024K used to pulldown full length PARD3 (A) or Dvl (B) ectopically expressed in HEK293T cells. **C)** GST-tagged PDZ domain of PARD3, Dvl, or mPDZ (PDZ3) used to pulldown ectopically expressed full length Daple-WT, E2024A, or E2024K in HEK293T cell lysates.

#### Supplementary References

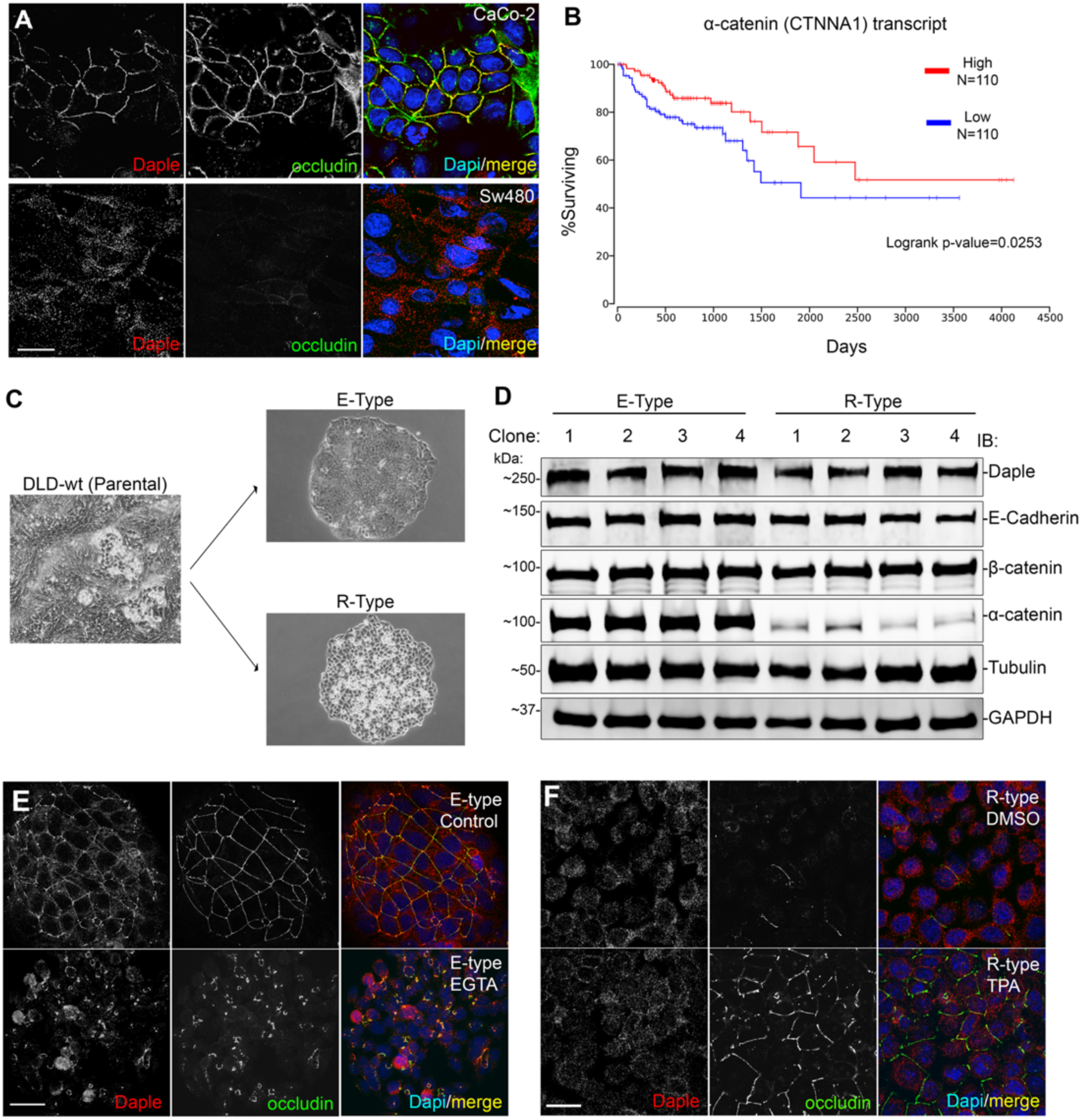


**Figure S1-Related to Figure 2. Daple localization and expression in colorectal cancer cell lines.**

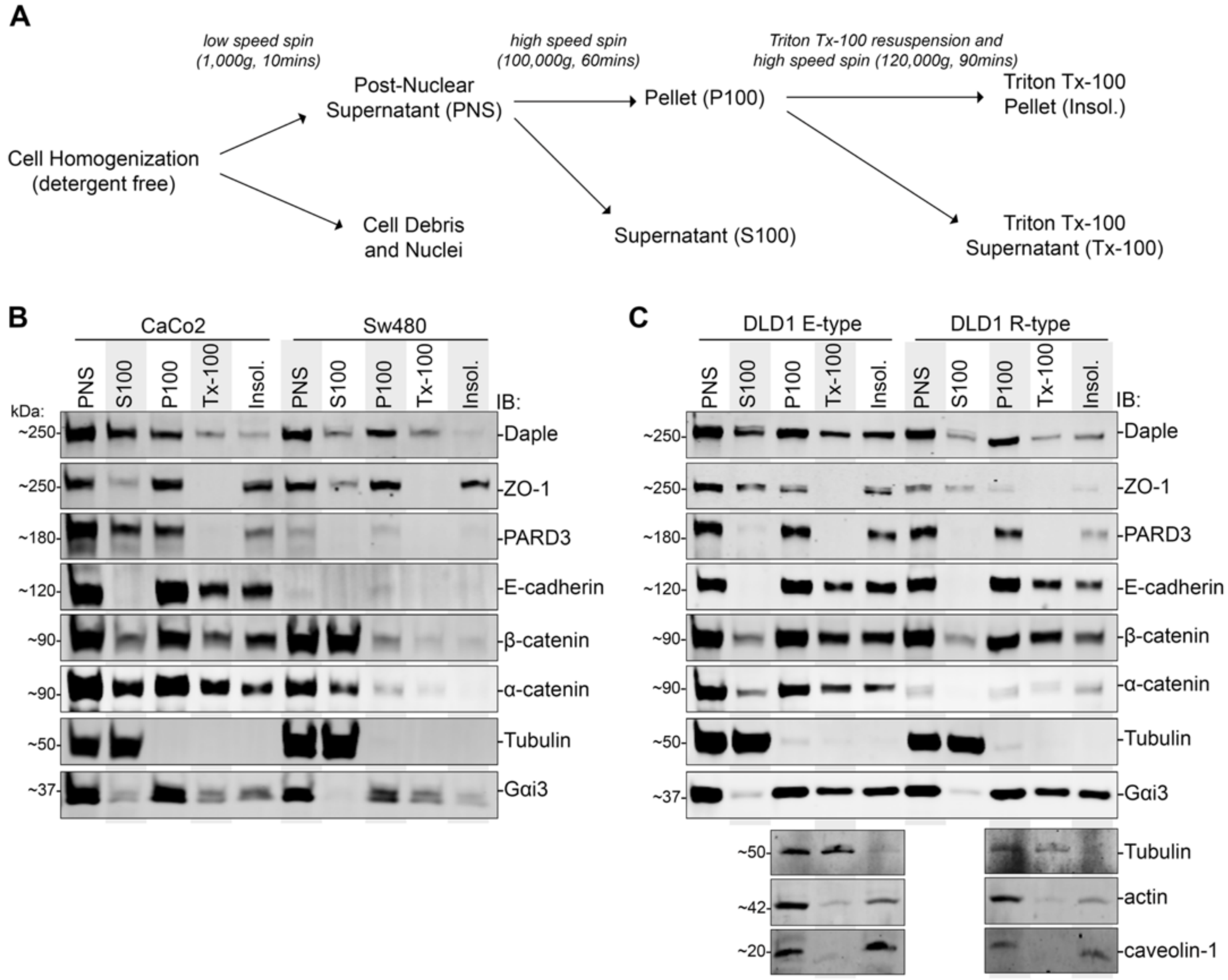


**Figure S2-Related to Figure 2. Subcellular fractionation studies on colorectal cancer cells reveal a pool of Daple that is on membrane and is detergent insoluble.**

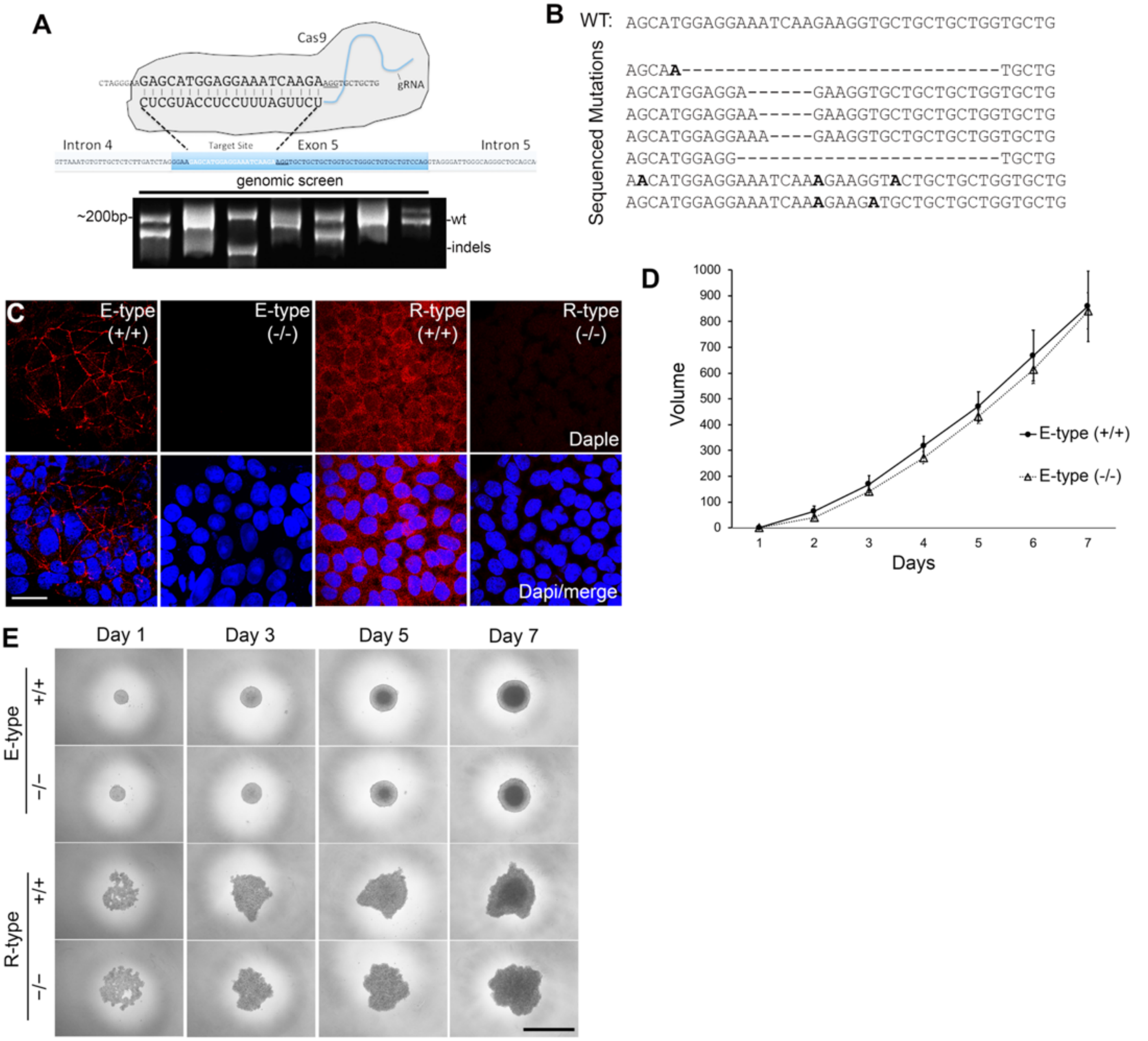


### Figure S3-Related to Figure 2. Characterization and validation of Daple CRISPR/Cas9 knockout

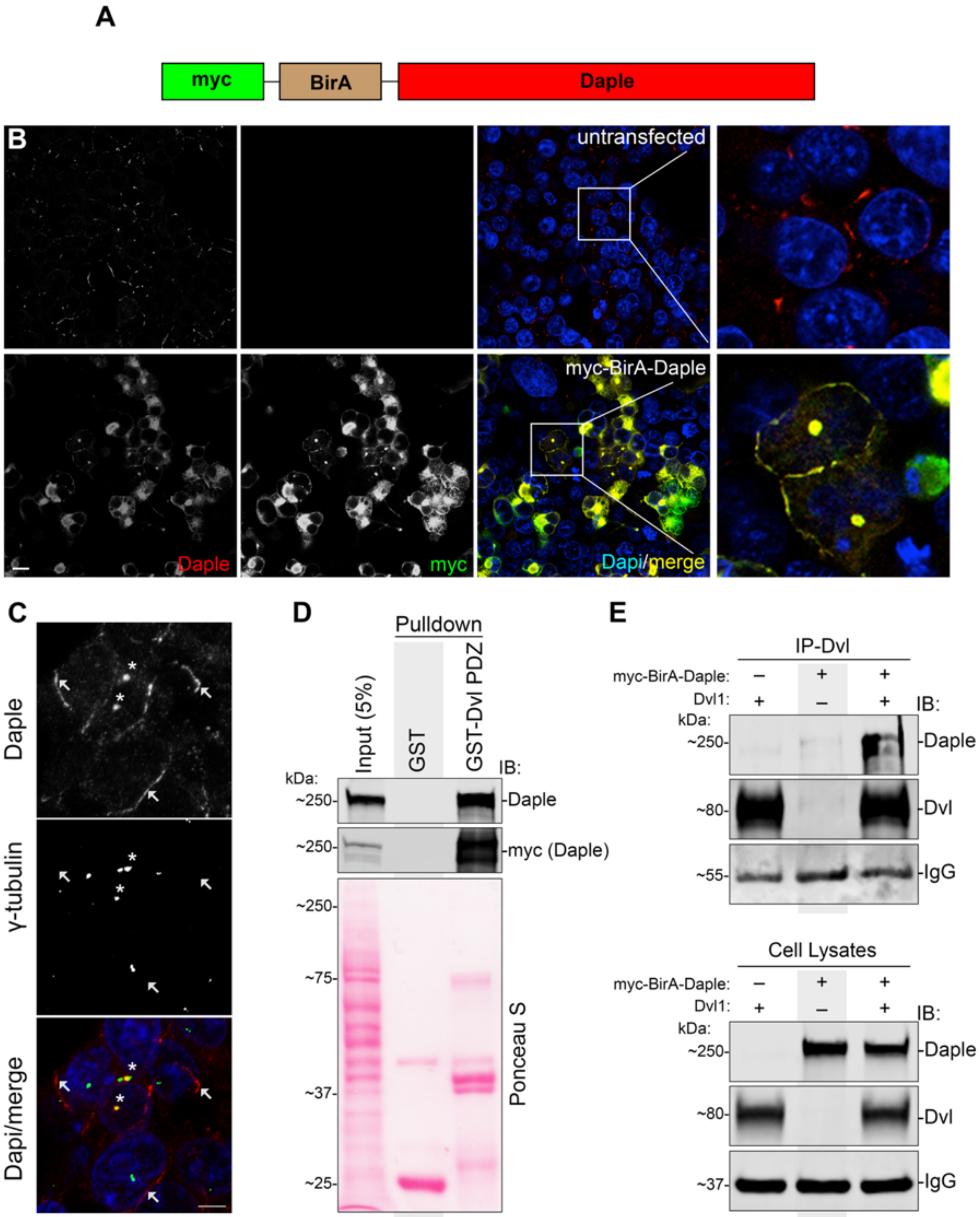


**Figure S4-Related to Figure 4. Validation of the Daple construct that was used in biotin proximity labeling.**

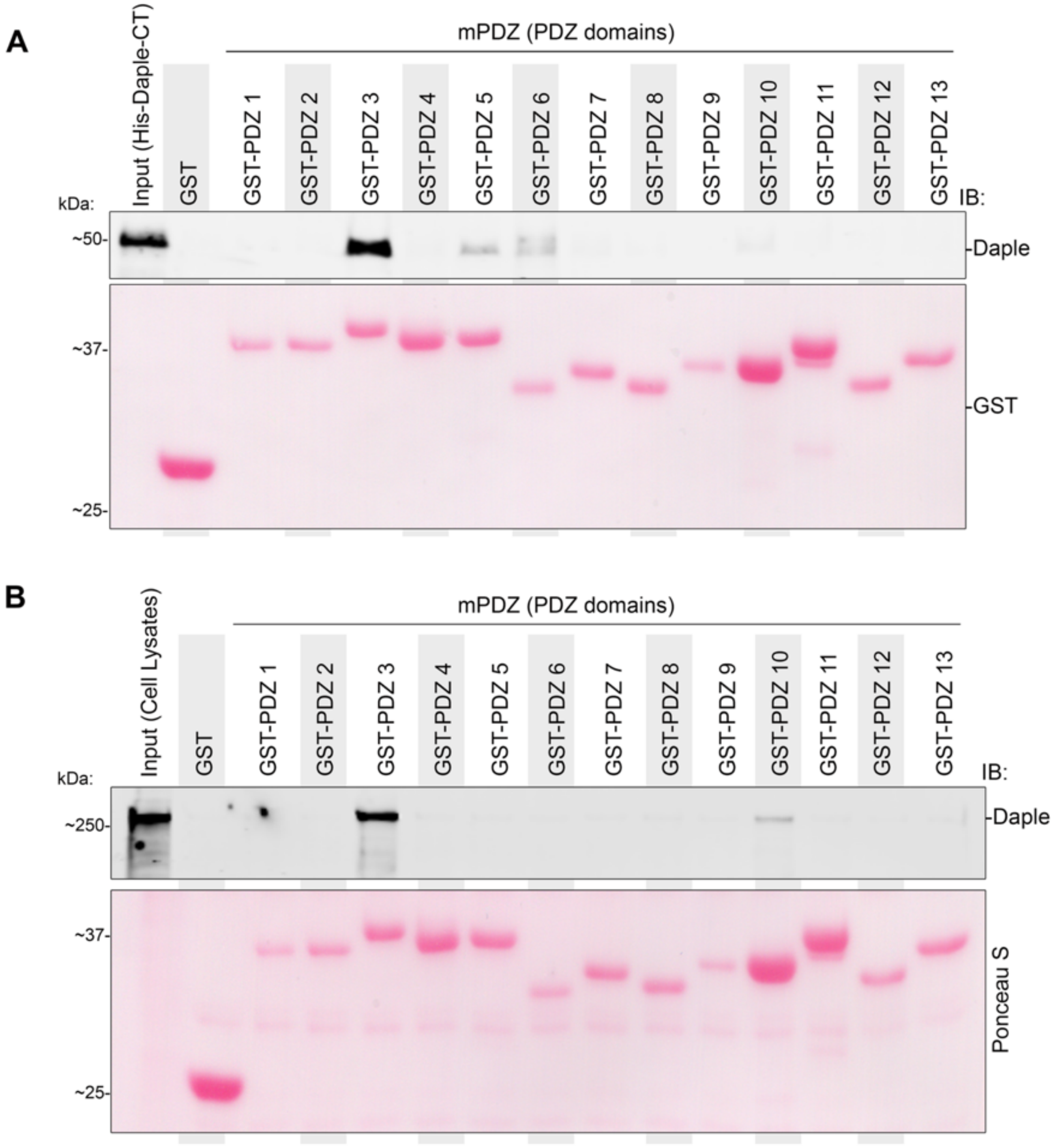


**Figure S5-Related to Figure 5. Daple binds directly to mPDZ; the interaction is mediated via Daple's PBM and the third PDZ module on mPDZ.**

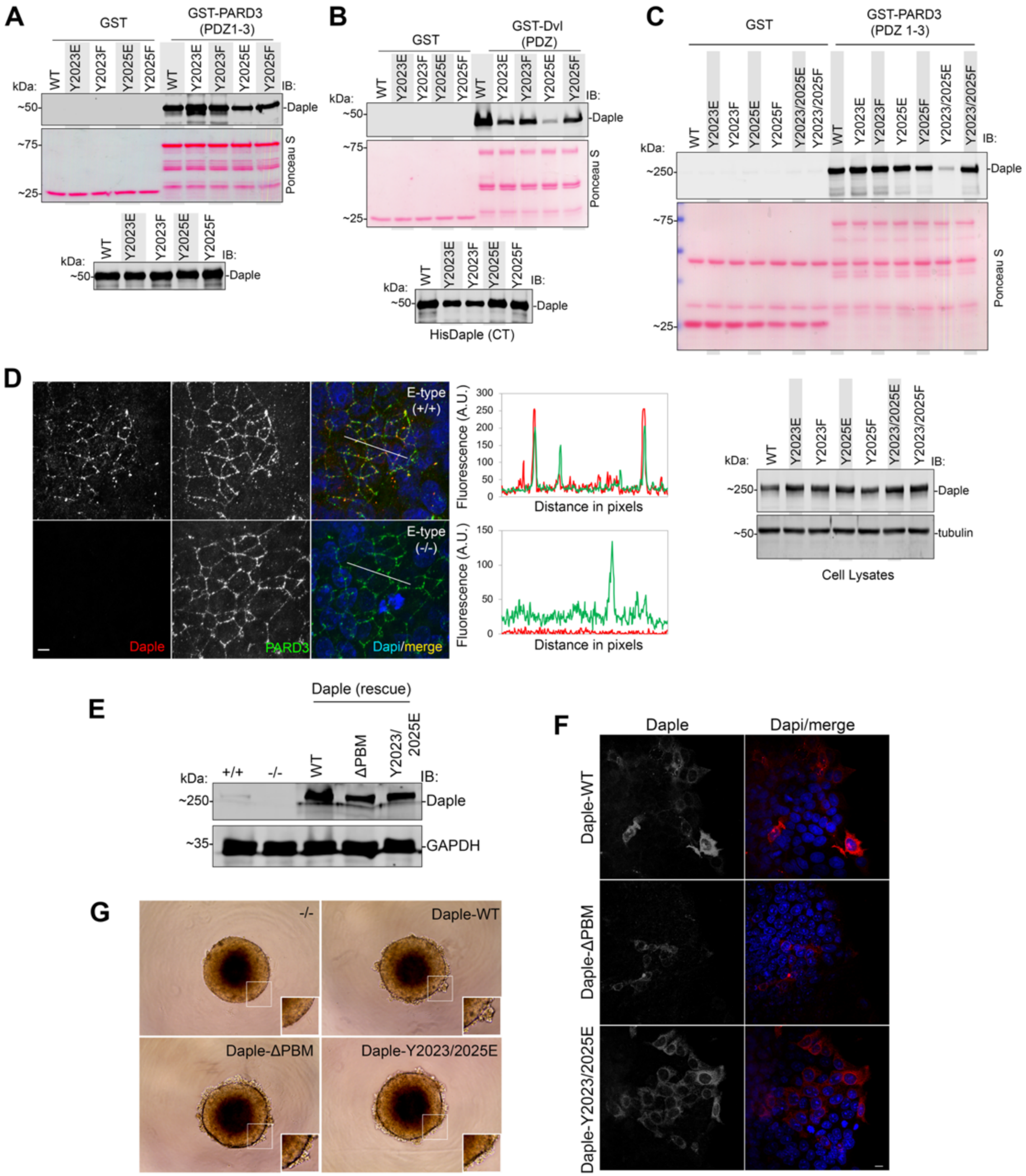


**Figure S6-Related to Figure 6. Impact of Daple and its tyrosine phosphorylation on binding and localization of PARD3.**

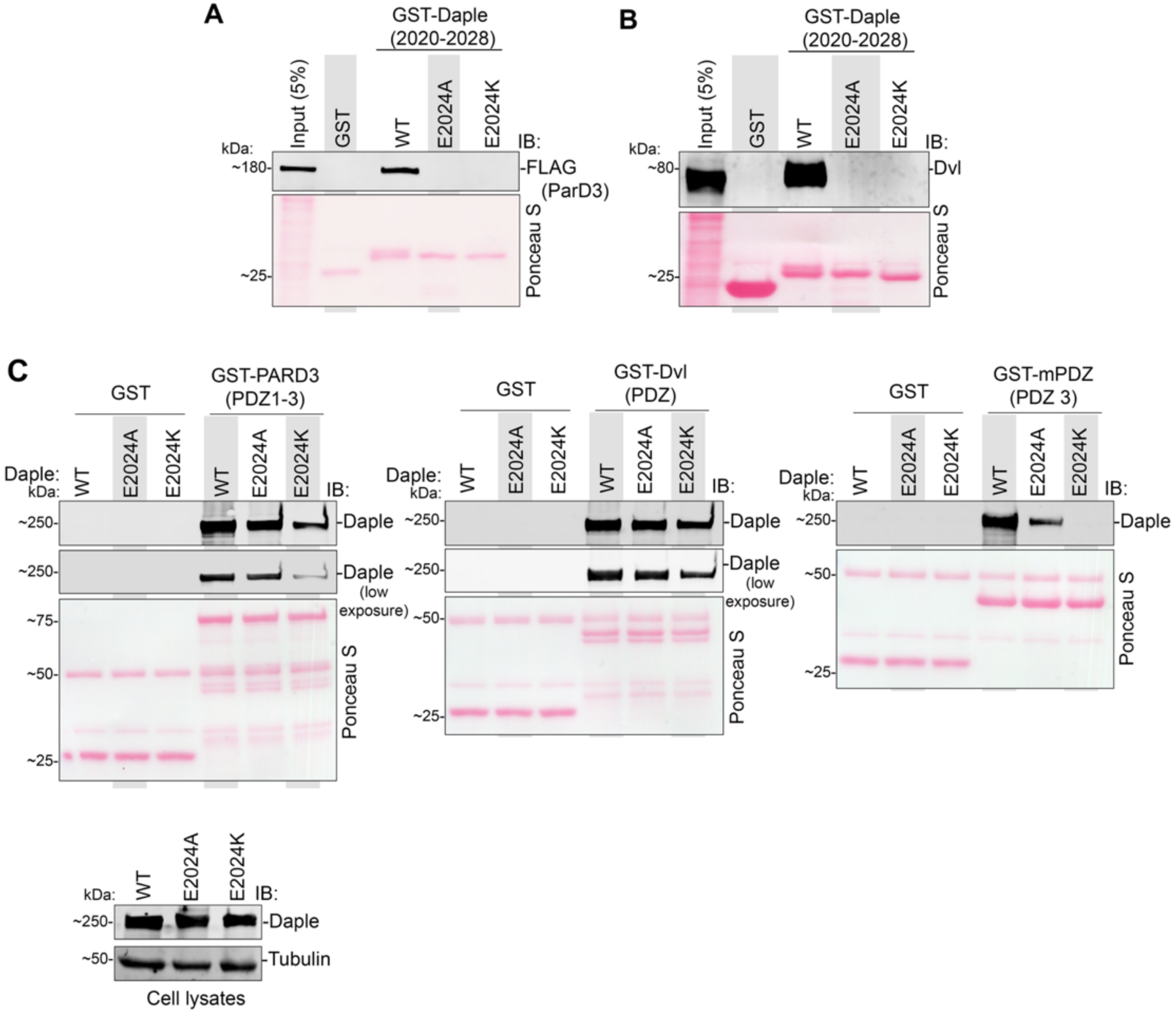


**Figure S7-Related to Figure 7. Impact of E2024 on Daple's ability to bind PDZ-domain containing proteins.**
